# MicroRNA-574 Regulates FAM210A Expression and Influences Pathological Cardiac Remodeling

**DOI:** 10.1101/2020.01.09.900712

**Authors:** Jiangbin Wu, Kadiam C Venkata Subbaiah, Feng Jiang, Omar Hadaya, Amy Mohan, Tingting Yang, Kevin Welle, Sina Ghaemmaghami, Wai Hong Wilson Tang, Eric Small, Chen Yan, Peng Yao

## Abstract

Aberrant synthesis of mitochondrial proteins impairs cardiac function and causes heart disease. However, the mechanism of regulation of mitochondria encoded protein expression during cardiac disease remains underexplored. Here, we have shown that multiple pathogenic cardiac stressors induce the expression of miR-574 guide and passenger strands (miR-574-5p/3p) in both humans and mice. miR-574 knockout mice exhibit severe cardiac disorder under heart disease-triggering stresses. miR-574-5p/3p mimics that are delivered systematically using nanoparticles reduce cardiac pathogenesis under disease insults. Transcriptome analysis of miR-574-null hearts uncovers *FAM210A* as a common target mRNA for both strands of miR-574. The interactome capture and translational state analyses suggest that FAM210A interacts with mitochondrial translation factors and regulates the protein expression of mitochondrial encoded electron transport chain genes. Using a human cardiomyocyte cell culture system, we discover that miR-574 regulates FAM210A expression and modulates mitochondrial encoded protein expression, which influences cardiac remodeling in heart failure.

## Introduction

Heart failure (HF) is a leading cause of morbidity and mortality worldwide^1^. The current 5-year mortality following diagnosis with HF is 52.6%^2^. An important cause of HF is pathological cardiac remodeling, which can be triggered by hypertension, coronary artery disease, chemotherapy, myocardial infarction (MI), valvular disease, or rare damaging genetic mutations. Following infarction and pressure overload, hypertrophic growth of cardiomyocytes (CMs), along with CM apoptosis and cardiac fibroblast (CF) proliferation and cardiac fibrosis, pathologically remodels the size, mass, geometry, and function of a heart, leading to HF. This pathological remodeling process is accompanied by changes in the expression of mitochondrial genes and specific miRNAs^3,4,5,6^.

The heart is the most mitochondria-rich organ in mammals. Mitochondria play critical roles in metabolism, cell proliferation, apoptosis, and CM contractility. The electron transport chain (ETC) complex, located in the mitochondrial inner membrane, generates an electrochemical-proton gradient that drives the synthesis of ATP. The synthesis of mitochondrial ETC complex proteins are tightly regulated in order to maintain mitochondrial homeostasis and normal cardiac function^7^. Accordingly, aberrant synthesis of mitochondrial proteins impairs heart function and causes heart disease. Early-stage cardiac hypertrophy induces mitochondrial protein translation and activity for a compensatory response, which may promote pathological cardiac remodeling^8^. In contrast, endpoint HF is accompanied by reduced mitochondrial protein synthesis and mitochondrial dysfunction^9^. Thus the restoration of mitochondrial function by targeting biogenesis pathways of mitochondrial proteins is thought to be an important strategy to develop therapeutics for heart disease^9,10,11,12^. However, this approach requires a better understanding of the regulatory mechanisms of mitochondrial genes. ETC complex genes contain mitochondrial-encoded (MEGs) and nuclear-encoded mitochondrial genes (NEMGs). Many studies have examined mechanisms underlying transcriptional regulation of these genes^13,14,15,16,17,18,19,20^. However, the role of regulation of protein expression of NEMGs and MEGs in cardiac disease has received little attention until recently. It has been shown that NEMG protein expression was enhanced in the murine heart with pressure overload^8^. Despite this finding, the regulation of MEG protein expression and the consequent coupling with NEMG protein expression in the heart remain largely underexplored.

miRNAs are 18-25-nt small non-coding RNAs that regulate gene expression via mRNA decay and/or translational repression. miRNAs are associated with Argonaute (Ago) proteins to form RNA-induced silencing complex (RISC), and guide miRISCs to specific mRNA targets^21^. Global loss of miRNAs in the murine heart via heart-specific deletion of Dicer produced a poorly developed ventricular myocardium, highlighting the importance of miRNAs in the heart^22^. Furthermore, abnormal expression of miRNAs has been observed frequently in response to pathological stress that impedes CM function and causes cardiac hypertrophy and HF^3, 23, 24^. Some miRNAs were explored as therapeutic targets to prevent HF^25,26,27,28^, while other miRNAs have been found to protect hearts from cardiac pathogenesis^23, 29, 30^. Inspired by all of these findings, stable miRNA mimics and antagonists for miRNAs have been developed to prevent or reverse various heart diseases in experimental HF mouse models^5^ and human clinical trials (e.g., miR-29 mimics and miR-92 inhibitors, miRagen Therapeutics, inc.). Previous studies have shown that miRNAs can regulate the translation of MEG mRNAs to affect heart function. For instance, mitochondrial-localized miR-1^31^ and miR-21^32^ directly target a selective cohort of seed sequence-specific MEG mRNAs and respectively activate their translation in skeletal muscle cells and cardiac myocytes. However, whether miRNAs also act through the regulation of general MEG protein expression to influence heart function is unclear.

In this study, we have discovered that miR-574 regulates FAM210A expression and antagonizes pathological cardiac remodeling. miR-574 null mice exhibit severe cardiac dysfunction under stress conditions, and exogeneous delivered miR-574 mimics protect against pathogenesis. Mechanistically, FAM210A functions as a novel regulatory factor of mitochondrial-encoded protein expression. Both the guide strand miR-574-5p and the passenger strand miR-574-3p of miR-574 target FAM210A to modulate the synthesis of mitochondrial-encoded ETC proteins. This novel mechanism may contribute to the maintenance of mitochondrial homeostasis and prevents pathological cardiac remodeling.

## Results

### Cardiac stress induces miR-574-5p and miR-574-3p in human and mouse hearts

From data mining of unbiased screening data from four laboratories, the guide strand miR-574-5p has been identified as a miRNA that is robustly induced in the heart in response to pathological cardiac remodeling, which included the cardiac tissues of patients during the initial stages following MI onset^33^, a mouse model of left coronary artery occlusion-derived MI^24^, and aged murine hearts (Fig. 1a)^26^. On the other hand, the passenger strand miR-574-3p is increased in the human hearts after MI and more remarkably induced by exercise training in murine hearts^34^. The cardiac functions of complementary strands of miR-574-5p and miR-574-3p remain unknown. miR-574 is conserved in 43 animal species (10 primates and 33 mammals, UCSC Genome Browser). In mammals, miR-574 is located in intron 1 of the host gene FAM114A1 (Family with sequence similarity 114 member A1) (Fig. 1b). We confirmed that both miR-574-5p and miR-574-3p were significantly induced in heart tissues of chronic HF patients, with the expression level of miR-574-5p even higher than that of miR-574-3p (Fig. 1c). Northern blot analysis showed that both miR-574-5p and miR-574-3p were expressed in the normal murine heart (Supplementary Fig. 1a). miR-574-5p and miR-574-3p were both significantly induced in isoproterenol (ISO)- or transverse aortic constriction (TAC)-treated mouse hearts, compared to vehicle treatment or sham operation (Fig. 1d and 1e). Moreover, in primary adult murine cardiomyocytes (ACMs), both strands of miR-574 were robustly induced by ISO treatment, with the guide strand showing even higher induction than the passenger strand, which is consistent with the results in chronic HF patients (Supplementary Fig. 1b).

**Fig. 1.**
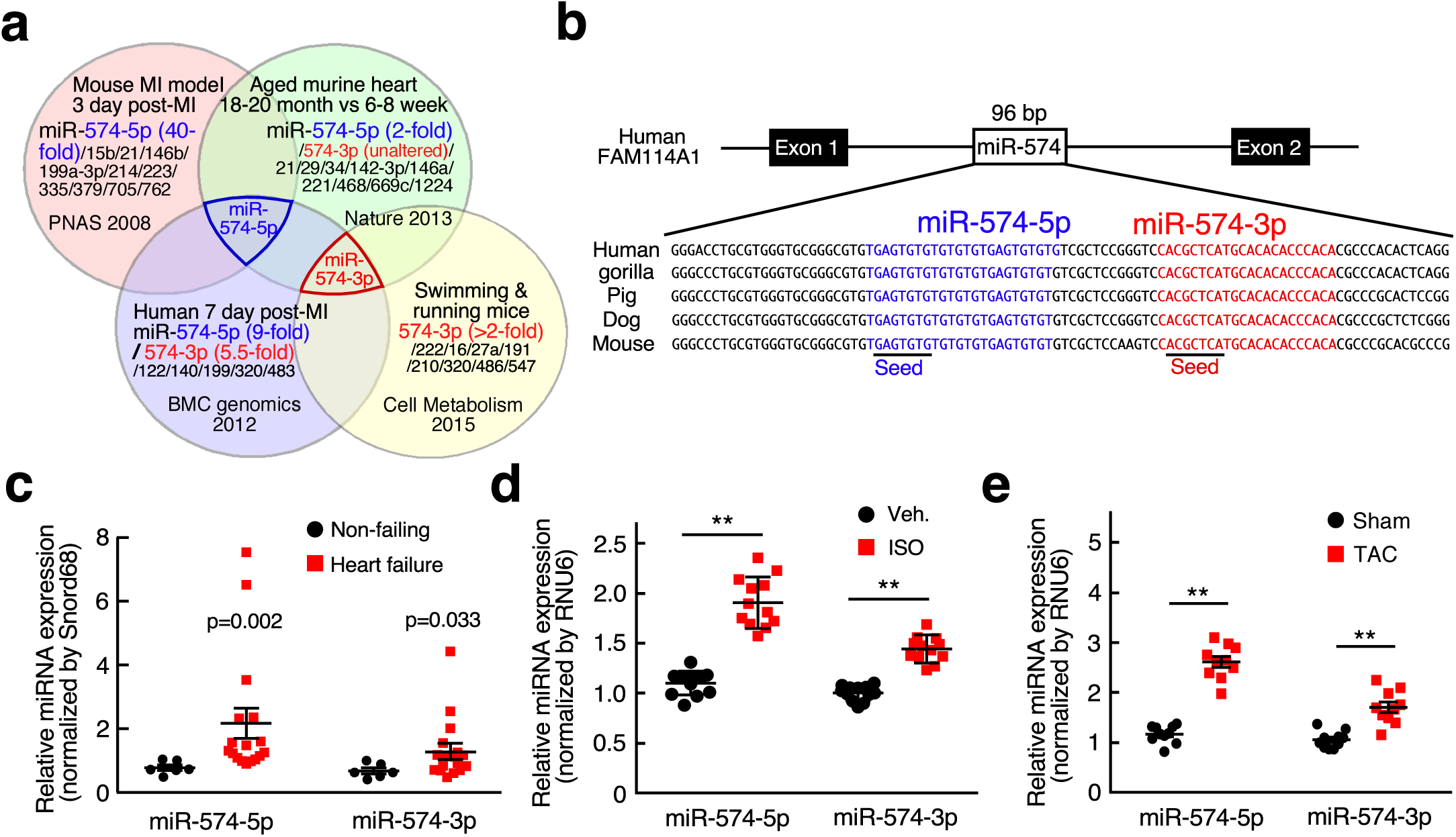
miR-574-5p and miR-574-3p expression is induced in human and mouse failing hearts. **a** miRNA profiling identified miR-574-5p as the commonly upregulated miRNA in the hearts of aged mice, MI mice, and patients; and miR-574-3p is induced in exercised hearts and MI patients. **b** Sequence and location of miR-574 gene in the human genome. **c** miR-574-5p and miR-574-3p were highly expressed in the explanted failing hearts from chronic HF patients (n=17) versus donor non-failing hearts (n=6). **d** miR-574-5p and miR-574-3p were induced in the hearts of mice with isoproterenol (ISO) infusion (n=10-12). **e** miR-574-5p and miR-574-3p were induced in the hearts of mice under transverse aortic constriction (TAC) surgery (n=8-10). Data were presented as mean ± SEM. *: p≤0.05, **: p≤0.01 by Student *t* test.

### miR-574 gene deletion exacerbates pathological cardiac remodeling

To determine whether miR-574 is protective or harmful in the adult murine heart, we generated a miR-574 knockout (KO) mouse using the embryonic stem (ES) cell clone harboring targeted miR-574 deletion (Supplementary Fig. 2a)^35^. The allele contained two loxP sites flanking the puromycin resistant cassette that replaces miR-574. Gene targeting in the ES cells was confirmed by qPCR (Supplementary Fig. 2b). The complete deletion of the genomic region between the two loxP sites was confirmed by PCR of genomic DNA (Supplementary Fig. 2c). Using RT-qPCR of RNA from adult murine hearts, we confirmed that the expression of miR-574-5p and miR-574-3p was abolished in the homozygous miR-574 KO mice (Supplementary Fig. 2d). Quantification of the *Fam114a1* mRNA level in miR-574^−/−^ hearts and isolated ACMs showed no change when probes spanning exon 1 and 2 were used (Supplementary Fig. 2e). miR-574^−/−^ mice were viable and appeared normal in weight, fertility, and behavior. We conclude that miR-574 is not required for viability and development in mice in the absence of stress.

We then investigated whether miR-574 affects cardiac remodeling in response to chronic β-adrenergic stimulation. Isoproterenol (ISO), a *β*-adrenergic receptor agonist, was administered to mice via subcutaneous injection (30 mg/Kg/day) for 4 weeks. The hearts of WT and miR-574^−/−^ mice were similar in size at baseline (Fig. 2a and Supplementary Fig. 2f). However, mutant mice showed a more significant ISO-induced increase in the left ventricle (LV) wall thickness and heart weight/tibia length (HW/TL) ratio, compared to WT mice (Fig. 2a, b). miR-574^−/−^ mice exhibited an enhanced cardiac hypertrophy phenotype associated with more enlarged CMs compared to WT mice after ISO treatment (Fig. 2c). Picrosirius red staining confirmed that ISO administration resulted in more fibrosis in miR-574^−/−^ than in WT mice (Fig. 2d). A hallmark of pathological hypertrophy and HF is the reactivation of a set of fetal cardiac genes. The cardiac gene activation was detected by RT-qPCR in WT and miR-574^−/−^ mice with vehicle or ISO treatment. Expression of the hypertrophic gene markers *ANF*, *BNP*, *Myh6*, and *Myh7*, and the fibrosis markers *Col1a2* and *Col3a1* were measured in the LV from the mice. Baseline expression of fetal cardiac genes was not altered in miR-574^−/−^ mice (Supplementary Fig. 2g). However, the expression of these genes was significantly increased in miR-574^−/−^ versus WT mice after ISO treatment (Fig. 2e), suggesting that miR-574^−/−^ mice are more susceptible to cardiac remodeling than WT mice in response to *β*-adrenergic stimulation. Primary ACMs from miR-574^−/−^ mice exhibited more severe ISO-induced hypertrophy (Fig. 2f) and were prone to excessive ISO-induced apoptosis (Fig. 2g, and Supplementary Fig. 2h) compared to those from WT mice. miR-574-5p is more dominantly expressed in human and mouse hearts under disease conditions (Fig. 1) and in isolated primary ACMs in comparison with miR-574-3p (Supplementary Fig. 2i). Transfection of miR-574-5p mimics antagonized ISO-induced hypertrophy of human ventricular cardiomyocyte cell line AC16 (Fig. 2h) while transfection of anti-miR-574-5p inhibitors promoted ISO-induced AC16 cell hypertrophy (Fig. 2i). We also tested the effect of miR-574-3p in ISO-induced cardiomyocyte hypertrophy. Transfection of miR-574-3p mimics reduced ISO-induced AC16 myocyte hypertrophy while anti-miR-574-3p exaggerated the effect (Fig. 2h, i). These data suggest that both miR-574-5p and miR-574-3p function as anti-hypertrophy factors in CMs.

**Fig. 2.**
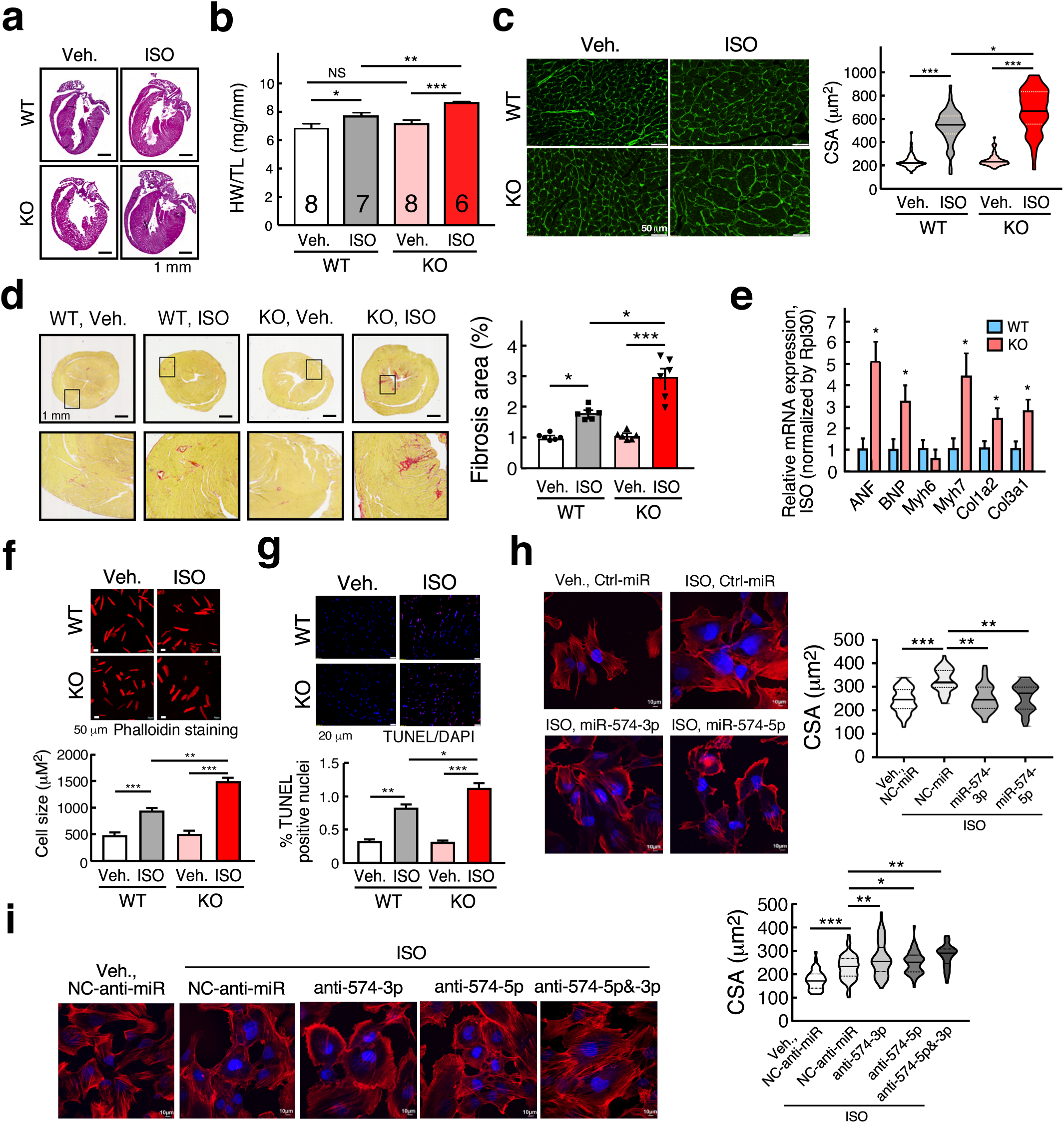
miR-574 gene knockout augments ISO-induced pathological cardiac remodeling. **a** H&E of hearts from WT and miR-574^−/−^ mice with or without ISO treatment for 4 weeks. Mice were between 8-10 weeks old females. **b** Ratio of HW/TL (heart weight/tibia length) in WT and miR-574^−/−^ mice (n=6-8 per group). **c** WGA (wheat germ agglutinin) staining in WT and miR-574^−/−^ mice. Cross-sectional area (CSA) of CMs was measured and quantified (n≥500 cells). **d** Picrosirius red staining in WT and miR-574^−/−^ mice. **e** RT-qPCR of fetal cardiac genes in WT and miR-574^−/−^ mice with ISO treatment. ANF, atrial natriuretic factor; BNP, B-type natriuretic peptide; Myh6/Myh7, myosin heavy polypeptide 6/7; Col1a2/Col3a1, procollagen, type I, *α*2; type III, *α*1. **f** Phalloidin staining of primary ACMs from WT and miR-574^−/−^ mice in response to ISO treatment (10 μM for 24 hrs). Data are obtained from 3 individual experiments (n=50). **g** TUNEL assay for heart tissue sections from WT and miR-574^−/−^ mice under ISO (10 μM for 48 hrs) versus vehicle treatment. **h** ISO-induced cardiomyocyte hypertrophy in human AC16 CM cells treated with miRNA mimics (100 nM for 24 hrs). n=50-100. **i** ISO-induced CM hypertrophy in AC16 cells treated with anti-miR inhibitors (100 nM for 24 hrs). n=50-100. Data were presented as mean ± SEM. *: p≤0.05, **: p≤0.01, ***: p≤0.001 by Student *t* test or one-way ANOVA.

To confirm the ISO-induced cardiac hypertrophic phenotype and the role of miR-574, we performed transverse aortic constriction (TAC) to model pressure overload-induced cardiac hypertrophy in WT and miR-574^−/−^ adult mice. miR-574^−/−^ mice exhibited more pronounced cardiac hypertrophy 4 weeks after initiation of TAC compared to WT mice, as indicated by HW/TL ratios (Fig. 3a, b) and CM cross-sectional area quantification (Fig. 3c). Cardiac fibrosis was also exaggerated in miR-574^−/−^ mice subjected to TAC, as demonstrated by picrosirius red staining (Fig. 3d). Consistent with the observed pathological changes, cardiac hypertrophy and fibrosis marker gene expression were increased more significantly in miR-574^−/−^ mice than in WT mice 4 weeks after TAC (Fig. 3e). Cardiac myocyte apoptosis was more pronounced in miR-574^−/−^ mice than in WT mice after TAC surgery (Supplementary Fig. 2j). Cardiac function of WT and miR-574^−/−^ mice in response to this cardiac stress was assessed by echocardiography, demonstrating a significantly impaired fractional shortening (FS) in miR-574^−/−^ compared to WT mice (Fig. 3f, and Supplementary Table 1). These studies suggest that miR-574^−/−^ hearts are prone to cardiac remodeling and functional deterioration in response to *β*-adrenergic stimulation and pressure overload, which implies the cardioprotective function of miR-574.

**Fig. 3.**
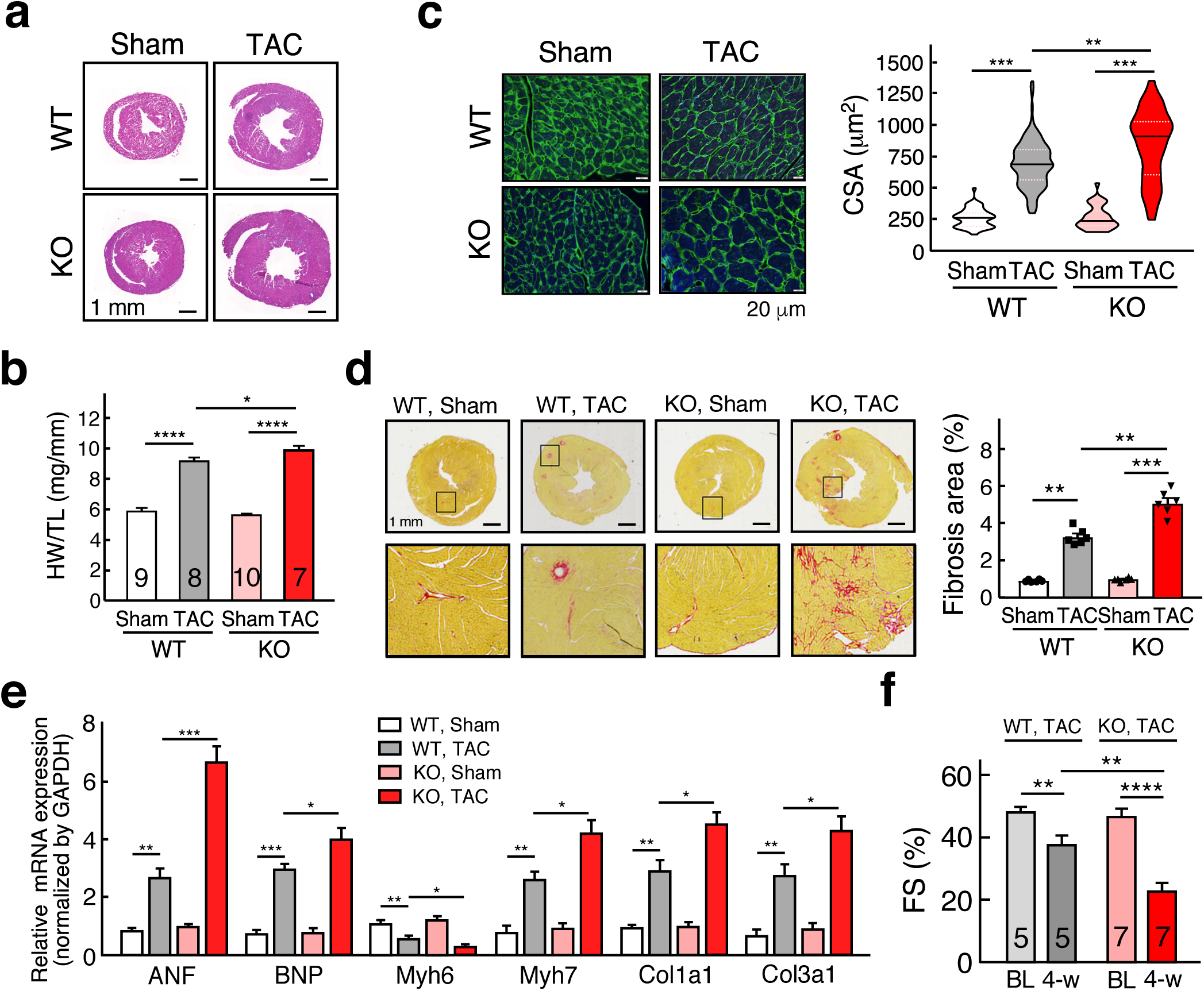
miR-574 gene knockout augments TAC-induced pathological cardiac remodeling. **a** H&E staining of hearts from WT and miR-574^−/−^ mice 4 weeks after TAC surgery. **b** Ratio of HW/TL in WT and miR-574^−/−^ mice (n=4-7 per group). **c** WGA staining of hearts from WT and miR-574^−/−^ mice 4 weeks after TAC surgery (n≥500 cells). **d** Picrosirius red staining of hearts from WT and miR-574^−/−^ mice under TAC surgery. **e** RT-qPCR of fetal cardiac genes in WT and miR-574^−/−^ mice with TAC surgery. **f** Echocardiography measurement of cardiac functions of hearts from WT and miR-574^−/−^ mice 4 weeks post TAC (n=5-7 per group). FS, fraction shortening. Data were presented as mean ± SEM. *: p≤0.05, **: p≤0.01, ***: p≤0.001 by one-way ANOVA.

### The therapeutic benefit of miR-574-5p/3p in cardiac remodeling using miRNA mimics

Having revealed a protective role of miR-574-5p and miR-574-3p in pathological cardiac remodeling, we sought to determine the therapeutic potential of these two miRNAs in the mouse heart *in vivo*. We used miRNA oligonucleotides that mimic endogenous miRNAs. We injected the mimetic oligomer of miR-574-5p, miR-574-3p, combined miR-574-5p/3p, or control miRNA (5 mg/Kg) into a tail vein of WT mice post TAC surgery once a week for 4 weeks, following an established strategy (Fig. 4a)^36^. *In vivo* miR-574-5p and miR-574-3p levels were measured in the heart to ensure efficient delivery and miRNA stability (Supplementary Fig. 3a). We first checked whether miR-574-5p and miR-574-3p target other organs, but did not observe any obvious adverse side-effects in the kidney and liver (Supplementary Fig. 3b). TAC-induced cardiac hypertrophy and fibrosis were significantly reduced by miR-574-3p mimics, as indicated by H&E and picrosirius red staining (Fig. 4a-c). miR-574-5p mimics were less effective compared to miR-574-3p, and a combination of miR-574-5p and miR-574-3p showed modest synergistic effects, possibly due to the limited loading efficiency of miR-574-5p/3p into the RISC complex in recipient cells. Expression of multiple hypertrophic and fibrotic marker genes was induced by the TAC surgery and reduced by miR-574-5p/3p treatment (Supplementary Fig. 3c). Cardiac function was partially recovered with miR-574-3p and miR-574-5p treatment (Fig. 4d and Supplementary Table 2). In addition, miR-574-5p/3p treatment reduced CM apoptosis after TAC surgery (Fig. 4e). We used a lower dose (1 mg/Kg) of miR-574-5p/3p mimics for injection, but we did not observe significant phenotypic improvement indicated by limited impact on cardiac hypertrophy and fibrosis (Supplementary Fig. 3e). Collectively, these results suggest that the phenotypic rescue is dependent on the sufficient dosage of miRNA mimics.

**Fig. 4.**
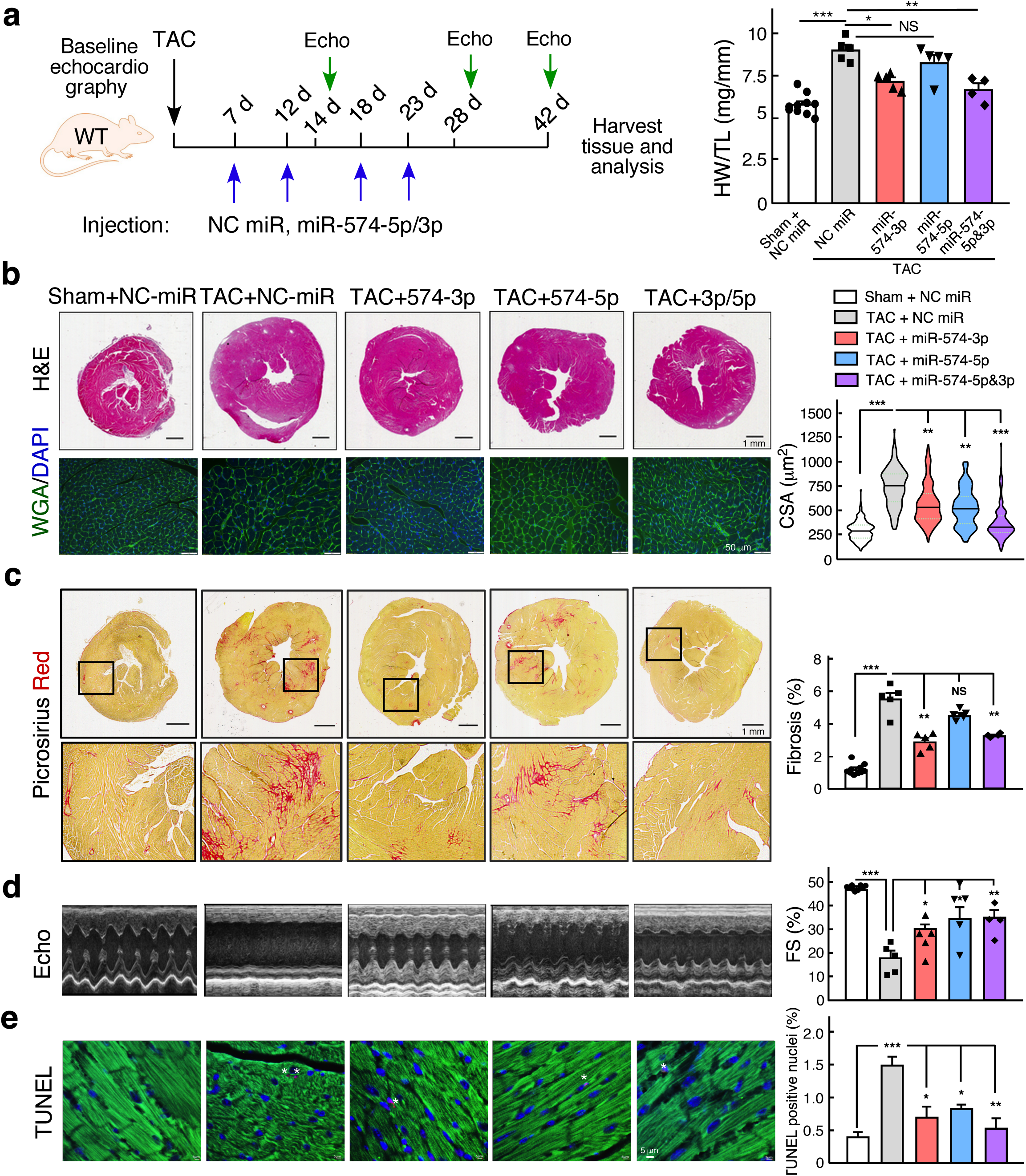
The therapeutic benefit of miR-574-5p/3p against cardiac pathological remodeling. **a** The schematic of therapeutic treatment of the TAC model using miRNA mimics followed by phenotypic characterizations. HW/TL ratio is measured (n=4-10 per group). NC, negative control. **b** H&E and WGA staining of murine hearts in the therapeutic models. **c** Picrosirius red staining of murine hearts in the therapeutic models. **d** Echocardiography analysis of the cardiac function of mice in the therapeutic models. **e** TUNEL assay of murine hearts in the therapeutic models. All the analyses were performed at the end-point of 6 weeks post-surgery. Data were presented as mean ± SEM. *: p≤0.05, **: p≤0.01, ***: p≤0.001 by one-way ANOVA.

### Transcriptome profiling identifies FAM210A as a common target of miR-574-5p/3p

To identify genes that are directly regulated by miR-574, we performed genome-wide RNA-Seq analyses with three pairs of P60 heart tissues from WT and miR-574^−/−^ mice at baseline. Protein-coding genes that are significantly altered in expression between miR-574^−/−^ and WT mice (*padj* < 0.05) were subject to Gene Ontology analyses. We identified 34 upregulated genes involved in the small molecule metabolism and the mitochondrial function, and 49 downregulated genes in the electron transport chain (ETC) complex, oxidative phosphorylation, and tricarboxylic acid cycle (Fig. 5a, b, Supplementary Fig. 4a, b, and Supplementary Table 3). The ETC genes (MEG: Nd1, Nd2, Nd3, Nd4, Nd5, Co1; NEMG: Ndufa12, Cox5b, Atpif1) and mitoribosome genes (mRpl32, mRps25) were reduced at modest but significant levels in miR-574^−/−^ versus WT hearts (Fig. 5b). Importantly, mitochondrial defects were not observed in the heart of miR-574^−/−^ mice at baseline, as indicated by ROS (reactive oxygen species) generation, membrane potentiality, ATP production, and mitochondrial morphology (Fig. 5c-e, and Supplementary Fig. 4c). These results imply that the reduction of ETC gene expression may be an adaptive response to maintain proper expression of these mitochondrial proteins. In contrast, in the heart of miR-574^−/−^ mice, ISO treatment triggered more pronounced ROS production (Fig. 5c), impaired membrane potentiality (Fig. 5d), compromised mitochondrial cristae formation (Supplementary Fig. 4c), and reduced ATP production (Fig. 5e).

**Fig. 5.**
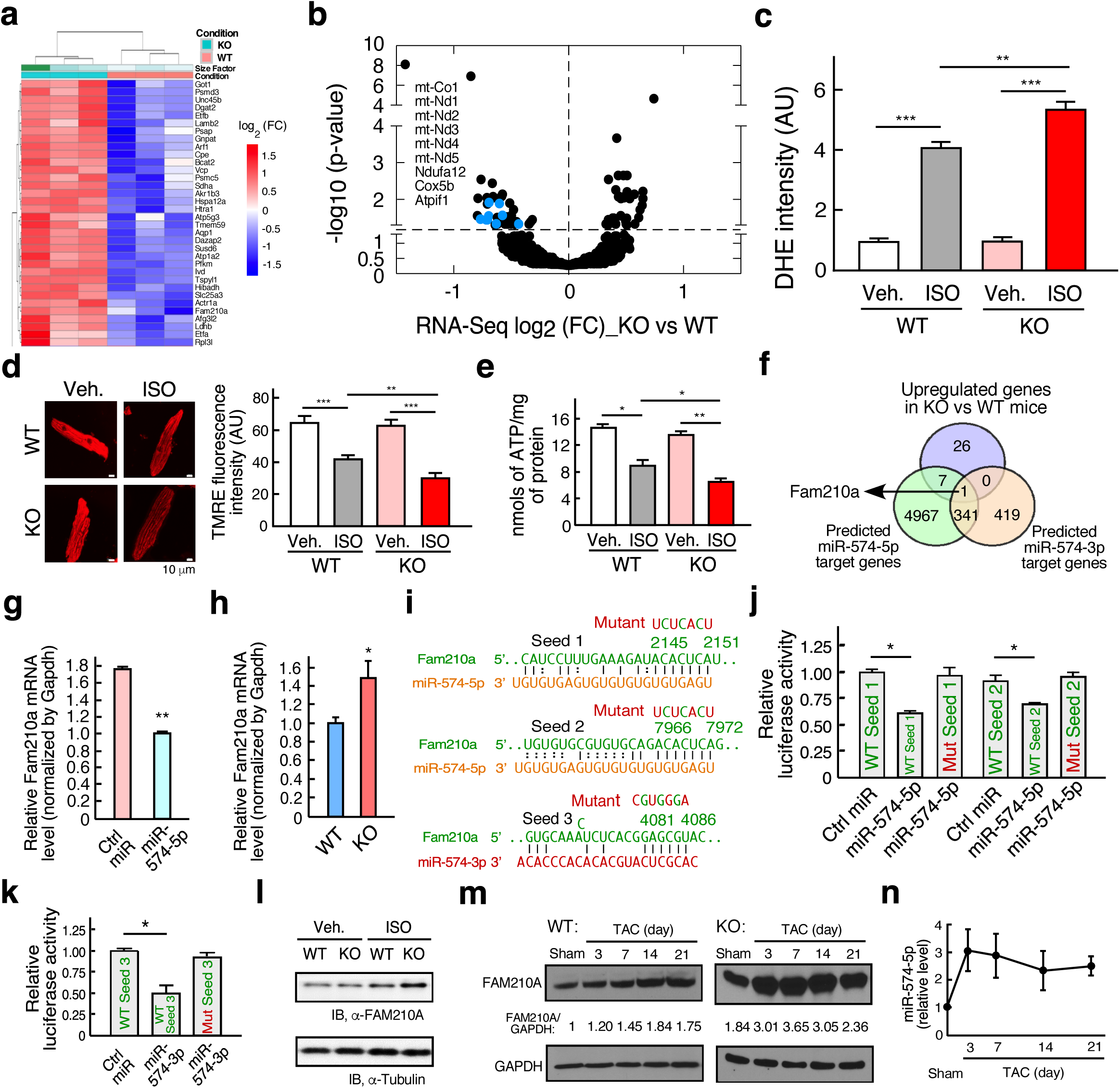
Identification of Fam210a as a common target of miR-574-5p/3p *in vivo*. **a** Heatmap of significantly upregulated genes in miR-574^−/−^ mice at baseline analyzed by RNA-Seq. P60 male mice, n=3 per group, padj<0.05. **b** Volcano curve analysis of dysregulated genes in miR-574^−/−^ versus WT heart. **c** DHE staining of isolated ACMs from WT and miR-574^−/−^ mice. **d** Mitochondrial membrane potentiality of ACMs from WT and miR-574^−/−^ mice. **e** ATP production in ACMs from WT and miR-574^−/−^ mice. **f** Identification of Fam210a as a target of miR-574-5p and miR-574-3p. Seed sequence-bearing target genes were predicted by TargetScan. **g** Validation of Fam210a as a miR-574-5p target using RT-qPCR analysis of mRNA expression. HL-1 CMs were transfected with control miRNA or miR-574-5p mimics. **h** Expression of *Fam210a* mRNA in the heart of miR-574^−/−^ and WT mice. **i** Seed sequences of miR-574-5p and miR-574-3p in *Fam210a* mRNA 3’UTR (TargetScan) and their mutants. **j** Dual luciferase assays with co-transfection of miR-574-5p mimics and FLuc-Fam210a 3’UTR bearing wild type and mutant seed sequences. **k** Dual luciferase assays with co-transfection of miR-574-3p mimics and FLuc-Fam210a 3’UTR bearing wild type and mutant seed sequences. **l** FAM210A protein expression from hearts of vehicle-or ISO-treated WT and miR-574^−/−^ mice (30 mg/Kg/day, s.c. injection for 4 weeks). **m** FAM210A protein expression during 21-day post-TAC in WT and miR-574^−/−^ mice. **n** miR-574-5p expression during 21-day post-TAC in WT mice. Data were presented as mean ± SEM. *: p≤0.05, **: p≤0.01, ***: p≤0.001 by Student *t* test or one-way ANOVA.

Eight of 34 upregulated genes were predicted to contain the target seed sequence of miR-574-5p (Fig. 5f). Seven out of these 8 genes are enriched in the small molecule metabolic pathway except for *Fam210a* (family with sequence similarity 210 member A), which is a novel gene with unknown function. Moreover, FAM210A is the only upregulated gene that contains the target seed sequence of both miR-574-5p and miR-574-3p in miR-574^−/−^ versus WT hearts. The seed sequences for both strands of miR-574 in *FAM210A* mRNA 3’UTR are present in mouse and human (Fig. 5f, and Supplementary Fig. 4d). Transfection of anti-miR-574-5p inhibitors increased expression of Fam210a and several other targets, such as LDHB and HIBADH (Supplementary Fig. 4e). Furthermore, the *Fam210a* transcript level was indeed increased in the heart of miR-574^−/−^ mice (Fig. 5h, and Supplementary Fig. 4f). *Fam210a* mRNA contains two target sites for miR-574-5p (Fig. 5i, top, middle). The dual luciferase reporter assay validated that each of the two wild type seed sequence sites of 3’UTR of *Fam210a* mRNA was a direct target of miR-574-5p. The inhibitory effect of miR-574-5p was abolished when we mutated the seed sequence in 3’UTR of the reporter constructs (Fig. 5i, j). *Fam210a* mRNA contains one target site of miR-574-3p (Fig. 5i, bottom), and the dual luciferase reporter assay confirmed that this miR-574-3p target site in *Fam210a* 3’UTR was functional (Fig. 5k). Upon ISO treatment, the protein expression of FAM210A was induced in the hearts of WT mice, and at a significantly higher level in miR-574^−/−^ mice (Fig. 5l). We measured the expression of miR-574-5p and FAM210A in WT and miR-574^−/−^ mice during a 21-day time course of TAC. In WT mice, the FAM210A protein level was not altered much at the early stage (day 3 and 7) but was significantly induced at the late stage (day 14 and 21) (Fig. 5m, left), while miR-574-5p was induced at both stages (Fig. 5n). In contrast, the FAM210A protein level was significantly induced at an early stage in miR-574^−/−^ mice (Fig. 5m, right). TAC surgery led to increased ROS production in the heart of miR-574^−/−^ mice compared to that of WT mice (Supplementary Fig. 5a). In addition, nanoparticle delivery of miR-574-5p and miR-574-3p mimics reduced FAM210A protein expression in the therapeutic mouse model (Supplementary Fig. 3d), which was accompanied with decreased ROS production and restored ATP production compared to control miRNA mimics injection in the mice under TAC surgery (Supplementary Fig. 5b, c). All these results suggest that FAM210A is a common target of miR-574-5p and miR-574-3p, and it may play a potential pathogenic role in cardiac remodeling.

### FAM210A binds to mitochondrial translation factors and regulates mitochondrial encoded protein expression

The function of FAM210A in the cardiac system and its molecular mechanism are entirely unknown. The FAM210A gene is conserved in 213 organisms, including human, dog, cow, mouse, rat, chicken, zebrafish, and frog (Supplementary Fig. 6a). RNA-Seq data showed that *Fam210a* mRNA is ubiquitously expressed in all organs, and is mostly enriched in the testis and heart (ventricle and atrium) of adult humans (Supplementary Fig. 6b, GTEx Portal)^37^. FAM210A mRNA and protein are expressed at higher levels in the hearts of cardiomyopathy patients compared to that of human non-failing hearts (Fig. 6a and Supplementary Fig. 6c). The expression of FAM210A protein was also increased in either ISO infusion or TAC surgery mouse models (Fig. 6b, c). Global homozygous Fam210a^−/−^ mice are embryonic lethal, while heterozygous Fam210a^+/–^ mice are viable and fertile^38^. LacZ reporter is highly expressed in the heart, brain, and skeletal muscle of Fam210a^+/–^ mice at E12.5 (Supplementary Fig. 6d, IMPC phenotyping data)^39^.

**Fig. 6.**
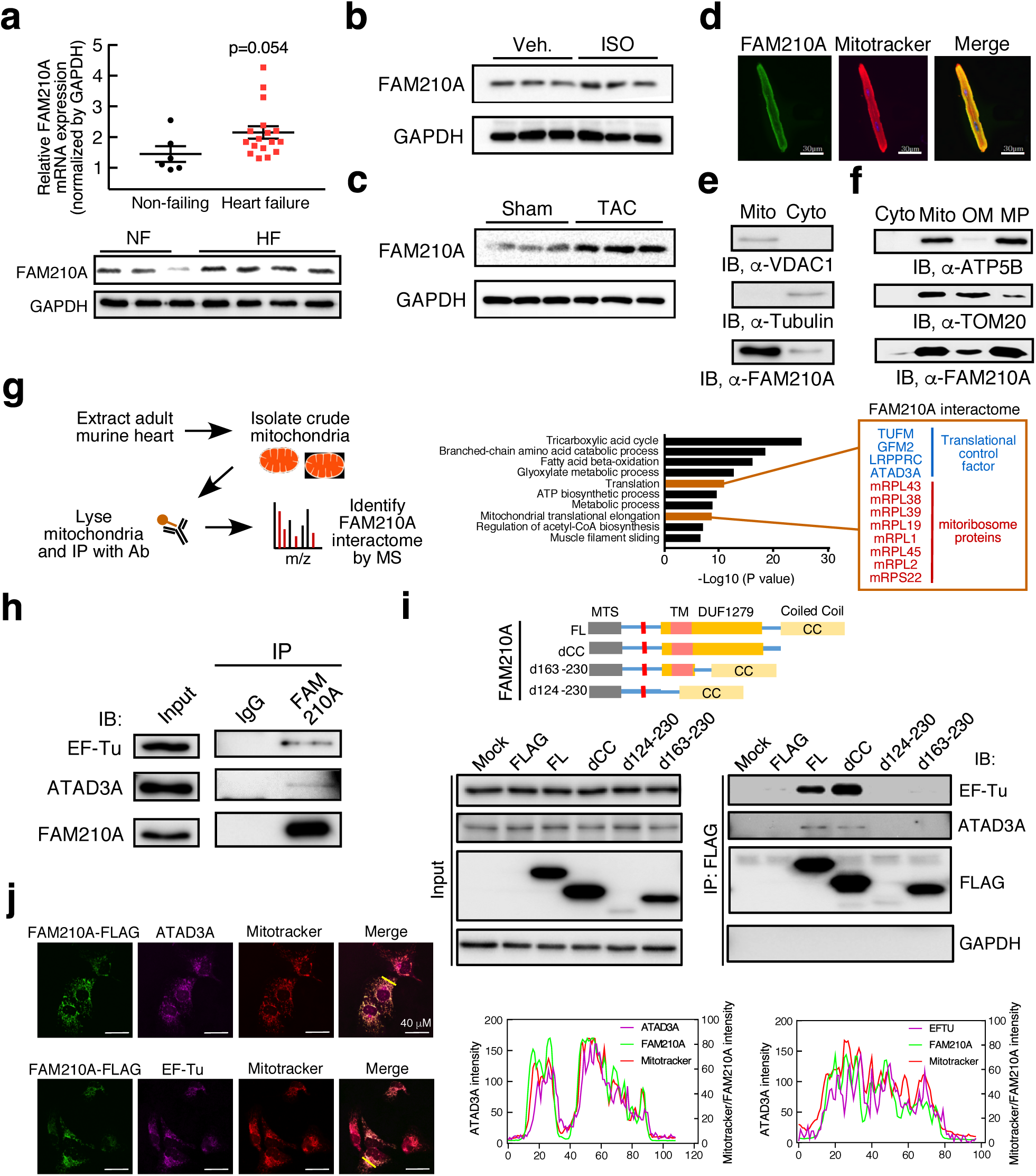
FAM210A is a mitochondrial protein, located in the inner membrane, and forms a complex with ATAD3A and EF-Tu. **a** RT-qPCR analysis of FAM210A expression in the explanted failing hearts from chronic HF patients (n=17) versus donor non-failing hearts (n=6). Western blot was done in a representative group of human samples. The statistical analysis is performed by Student *t* test. **b** The expression level of FAM210A protein in ISO-treated murine hearts (n=3, ISO minipump infusion, 20 mg/Kg/day, 4 weeks). **c** The expression level of FAM210A protein in TAC-treated murine hearts (n=3). **d** Cellular localization of FAM210A in primary mouse ACMs detected by IF. **e** Immunoblot detection of FAM210A in isolated cytoplasmic and mitochondrial fractions from mouse hearts. VDAC1 and tubulin are used as fraction controls. Cyto: cytoplasm; Mito: mitochondria. A representative image is shown from three biological replicates. **f** Sub-organelle localization of FAM210A in mitochondria. TOM20 and ATP5B are used as the outer membrane (OM) and mitoplast (MP) markers, respectively. **g** Antibody-based affinity purification of endogenous FAM210A from purified murine cardiac mitochondria coupled with mass spectrometry analysis. **h** IP-IB validates FAM210A-interacting proteins using pre-immune IgG and FAM210A antibodies. A representative image is shown in three biological replicates. **i** Mapping of interacting domains in FAM210A for binding EF-Tu and ATAD3A. **j** Confocal imaging of co-localization of FAM210A-3xFLAG and endogenous ATAD3A or EF-Tu by immunostaining.

FAM210A is a transmembrane protein localized in the mitochondria^40, 41^. FAM210A contains a mitochondrial targeting signal (MTS; cleavage site around Val^95^, MitoCarta 2.0) peptide, a DUF1279 (Domain of Unknown Function 1279) domain containing a transmembrane (TM) peptide (predicted by TOPCONS server^42^), and a coiled coil (CC) at C-terminus (EMBL-EBI bioinformatics) (Supplementary Fig. 6e). Immunofluorescence and cellular fractionation followed by immunoblot confirmed that endogenous and overexpressed FAM210A was mainly localized in the mitochondria of the mouse ACMs (Fig. 6d), murine heart (Fig. 6e), HEK293T human embryonic kidney cells (Supplementary Fig. 6f), and AC16 human CM cells (Supplementary Fig. 6g,h). Fractionation of mouse cardiac mitochondria showed that the majority of FAM210A was localized in mitoplasts (mitochondrial inner membrane and matrix), and a small fraction was located at the mitochondrial outer membrane (Fig. 6f). Prior mass spectrometry screens have reported that FAM210A is the strongest binding protein tightly associated with ATAD3A (ATPase family AAA domain-containing protein 3A) and mitoribosome^41^. Also, ATAD3A is associated with mitochondrial elongation factor EF-Tu (TUFM)^41^. ATAD3A is an essential factor for mitochondrial translation, and its genetic mutations cause Harel-Yoon syndrome with hypertrophic cardiomyopathy^43, 44^. To confirm the interaction between FAM210A and ATAD3A or EF-Tu, we used a FAM210A-specific antibody to pull down the endogenous FAM210A protein from the mitochondria purified from the mouse hearts for mass spectrometry analysis of its interactome (Fig. 6g). We identified the interactome of FAM210A, including a number of proteins in the translation machinery, such as translation factors and mitoribosome proteins, implying that it may participate in translational control (Supplementary Table 4). These interacting translation factors include TUFM/EF-Tu, GFM2 (mitochondrial ribosome releasing factor), ATAD3A, and LRPPRC. IP and IB for endogenous FAM210A confirmed EF-Tu and ATAD3A were bona fide interacting proteins (Fig. 6h). We further confirmed the interaction between FAM210A and EF-Tu or ATAD3A using FLAG-tagged FAM210A (Supplementary Fig. 6i). We mapped the interacting domain of FAM210A with EF-Tu and ATAD3A using a series of truncated FAM210A mutants (Fig. 6i). We found that the DUF1279 domain was responsible for binding EF-Tu and ATAD3A, but the C-terminal CC domain was not required for binding. Instead, the deletion of the CC domain enhances the interaction between FAM210A and EF-Tu or ATAD3A, suggesting that the CC domain might function as a regulatory module to influence the interaction with EF-Tu or ATAD3A. The deletion of the single TM domain led to deficient expression of the truncated protein, possibly due to mislocalization and subsequent degradation of the mutant FAM210A protein. To further confirm that FAM210A is associated with EF-Tu or ATAD3A in the mitochondria, we performed co-immunostaining for ectopically expressed FLAG-tagged FAM210A and endogenous EF-Tu and ATAD3A. Double immunostaining and two-color histogram analysis revealed significant co-localization between FAM210A and EF-Tu or ATAD3A in the mitochondria, respectively (Fig. 6j).

Since FAM210A interacts with mitochondrial translation factors and may play a general role in multiple cell types and organs^38^, we used the easy-to-transfect HEK293T cells to study its molecular mechanisms in gene regulation. To examine whether FAM210A affects transcription of mitochondrial genes, we performed the RNA-Seq after siRNA-mediated knockdown of FAM210A. The transcriptomic profiling data revealed that mitochondrial gene expression (including NEMG and MEG) was not dramatically altered at the mRNA level in the acute phase (Supplementary Fig. 6j, and Supplementary Table 5). This result suggests that FAM210A does not play a role in transcriptional control of mitochondrial genes. Next, we tested whether FAM210A regulates the translation of mitochondrial genes. We first performed polysome profiling of the cytosolic fraction upon FAM210A overexpression (Supplementary Fig. 7a, left). No change was observed in the global polysome profile state in the cytoplasm, implying that FAM210A does not regulate cytosolic translation (Supplementary Fig. 7b). Moreover, siRNA-mediated knockdown of FAM210A did not alter either global polysome profile or translation of NEMG mRNAs from all five ETC complexes (Supplementary Fig. 7c). We then tested the possibility that FAM210A is involved in the regulation of mitochondrial translation. The short time pulse-chase AHA labeling assay showed that overexpression of FAM210A could increase the translation of multiple MEG proteins (Supplementary Fig. 7d). These proteins include several mitochondrial-encoded ETC complex components from the Complex I, III, IV, and V. We then performed polysome profiling of the mitochondrial fraction of FAM210A overexpressed cells using mock transfection as control (Supplementary Fig. 7 a, right). qRT-PCR following polysome profiling showed that mRNAs of ND3, CYTB, CO1, and ATP6 were more enriched in the large mitoribosome subunit and the heavy polysome fractions (Supplementary Fig. 7e), suggesting more efficient uploading of the large ribosome subunit and accelerated translational elongation. This observation is consistent with the finding of an interaction between FAM210A and EF-Tu and ATAD3A, suggesting that FAM210A works with EF-Tu to increase translation efficiency in mitochondria at the elongation step. In summary, FAM210A modulates the translation efficiency of multiple MEGs encoding ETC component proteins at the translational elongation step, possibly by interacting with EF-Tu and ATAD3A (Supplementary Fig. 7f).

### miR-574 influences mitochondrial encoded protein expression via regulation of FAM210A

Previous studies have shown that NEMG mRNA translation is enhanced during TAC surgery using translating ribosome affinity purification (TRAP) coupled with RNA-Seq of the ribosome-bound mRNAs (Supplementary Fig. 7g)^8^. We hypothesized that FAM210A might contribute to the increase of MEG protein expression that may outmatch the surge in NEMG protein expression, while miR-574 limits expression of MEG proteins and maintain the MEG-NEMG protein homeostasis by downregulating FAM210A in mitochondria. To assess this hypothesis, we first measured the MEG and NEMG protein expression after overexpression or knockdown of FAM210A in human AC16 cardiomyocyte cells. We found that overexpressed or knockdown of FAM210A increased or decreased MEG protein expression, respectively, but not NEMG protein expression (Fig. 7a, b, Supplementary Fig. 8a, b), suggesting that FAM210A promotes the protein expression of MEGs but not NEMGs. We then measured the MEG and NEMG protein expression with transfection of miR-574-5p, miR-574-3p, or the anti-miR inhibitor of each. Overexpression of either miR-574-5p or miR-574-3p reduced MEG protein expression (Fig. 7c, Supplementary Fig. 8c), while transfection of anti-miR-574-5p or anti-miR-574-3p inhibitor increased MEG protein expression (Fig. 7d, Supplementary Fig. 8d). In contrast, in both experiments, NEMG protein expression was not affected. These observations suggest that miR-574 downregulates MEG protein expression by targeting FAM210A. Moreover, co-overexpression of FAM210A in the presence of miR-574-5p/3p mimics transfection rescued MEG expression (Fig. 7e, Supplementary Fig. 8e) without affecting NEMG protein expression, further confirming the role of miR-574-FAM210A axis in regulating MEG protein expression in human CMs. To examine whether the excessive expression of MEG proteins by FAM210A and normalization by miR-574-mediated inhibition affect mitochondrial activity, we measured the mitochondrial phenotypes in the same set of treated cells in the presence or absence of ISO stimulation. We found that overexpression of FAM210A reduced mitochondrial membrane potentiality and ATP production, and co-overexpressed miR-574-5p and miR-574-3p partially restored both (Fig. 7f, g). In addition, overexpression of FAM210A also exaggerated hypertrophic growth of myocyte cells, and co-overexpressed miR-574-5p and miR-574-3p normalized the cell size (Fig. 7h), which is consistent with partially restored mitochondrial functions (Fig. 7f, g). These observations in human cardiomyocyte cells suggest that the miR-574-FAM210A axis prevents excessive expression of MEG proteins in the ETC complex, thereby maintaining mitochondrial protein homeostasis and normal mitochondrial functions. This mechanism may contribute to the cardioprotective effects of miR-574 in the mouse HF models.

**Fig. 7.**
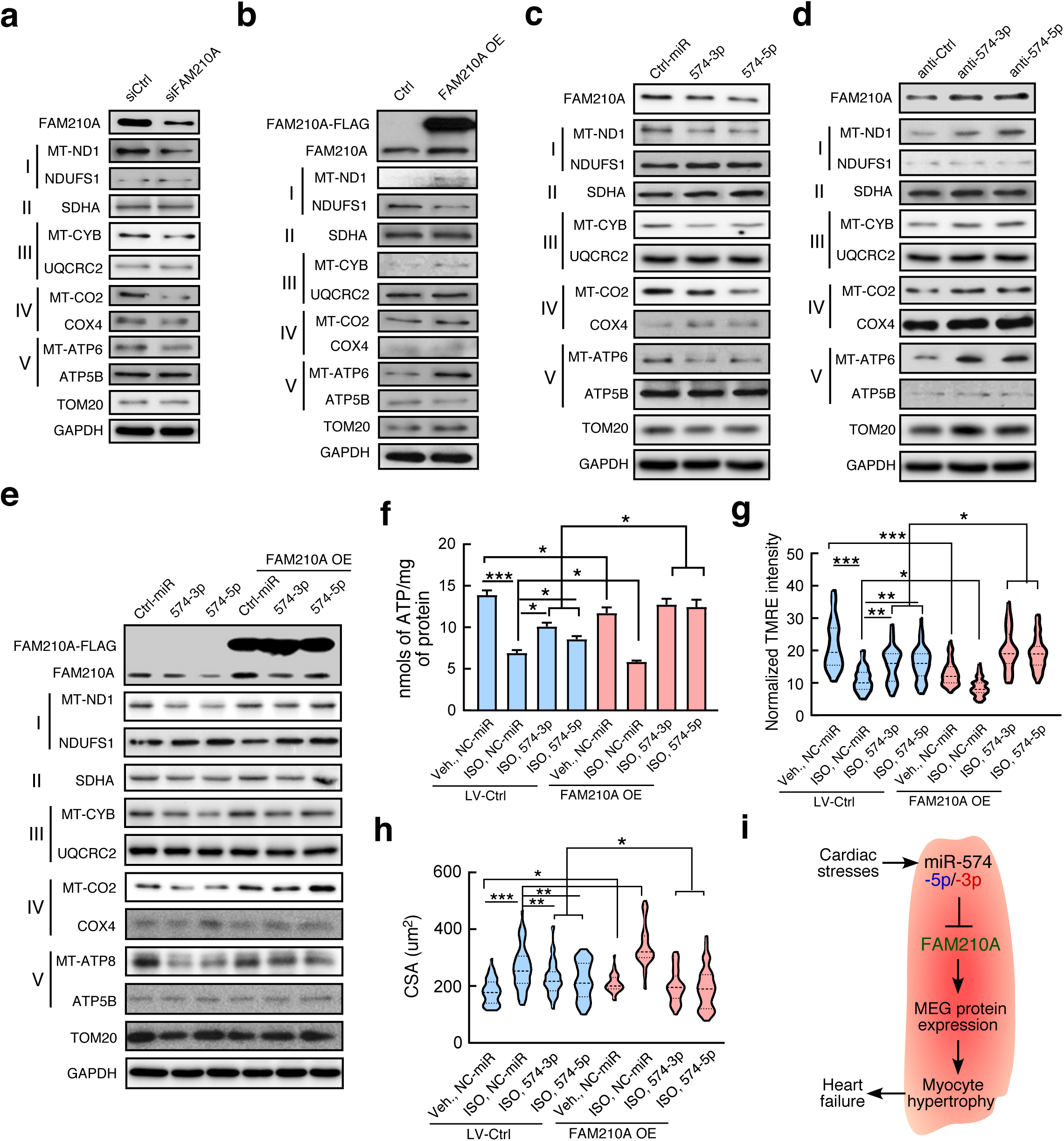
miR-574-FAM210A axis modulates the expression of mitochondrial proteins and mitochondrial activity. Protein expression of mitochondrial ETC component genes was measured in AC16 cells after following treatments in comparison with control cells: **a** siRNA knockdown of FAM210A. **b** FAM210A overexpression. **c** miR-574-5p or miR-574-3p overexpression. **d** Transfection of anti-miR-574-5p or anti-miR-574-3p inhibitor. **e** transfection of miR-574-5p/3p in the absence or presence of FAM210A overexpression. All the Western blot images were provided as representative data from 3-4 biologically replicated experiments. Quantitative data were shown in Fig. S8a-e. **f-h** ATP production, mitochondrial membrane potentiality, and CM hypertrophy in AC16 cells under various conditions. Data were presented as mean ± SEM. *: p≤0.05, **: p≤0.01, ***: p≤0.001 by Student t test or one-way ANOVA. **i** The schematic model of the cardioprotective function and mechanism of miR-574-FAM210A axis against cardiac pathological remodeling.

## Discussion

This study was initiated from our unbiased screening of heart disease-related miRNAs in multiple independent studies (Fig. 1a). We found that miR-574-5p and miR-574-3p were induced in human and mouse diseased hearts. We demonstrated that miR-574-5p/3p play cardioprotective roles using a miR-574 genetic knockout mouse model and injection of miR-574-5p/3p mimics in mouse HF models. RNA-Seq of baseline miR-574-null mouse hearts revealed the mRNA of *Fam210a* as a critical downstream target of both strands of miR-574. FAM210A functions directly or indirectly in regulating mitochondrial encoded protein expression. miR-574 restricts FAM210A expression and may act as a molecular brake to maintain mitochondrial homeostasis and normal cardiac function.

The guide and passenger strands of miR-574 are both loaded onto the RISC. We used synthesized miR-574-5p and miR-574-3p to make a standard curve based on RT-qPCR and measure the absolute copy number of each miRNA in CM versus CF. We found that miR-574-5p and miR-574-3p were both expressed at the moderate level in freshly isolated CM and CF (Supplementary Fig. 1c). miR-574-5p has been identified as an oncogenic miRNA^45^ and is involved in neurogenesis^46^. miR-574-3p is recognized as an anti-tumor miRNA^47, 48^. miR-574-5p/3p are differentially induced in the heart under various conditions. miR-574-5p displays dominant expression in CMs compared to miR-574-3p in human and mouse failing hearts as well as in aged mouse hearts (Fig. 1a). Recently, miR-574-5p has been shown to be dysregulated in two hypertrophic cardiomyopathy mouse models, including R92W-TnT and R403Q-*α*MyHC mutation-bearing mice^49^. In contrast, miR-574-3p is induced in the hearts of mice that undergo exercise training (Fig. 1a). Using a loss-of-function model, we show that miR-574^−/−^ mice display more severe cardiac hypertrophy and fibrosis compared to WT mice after ISO injection and TAC surgery, suggesting a cardioprotective function of miR-574. Both strands of miR-574 downregulate a common target gene, FAM210A, by binding to three different regions of its mRNA 3’UTR. Overexpression or inhibition of miR-574-5p (and also miR-574-3p) levels in AC16 human CM cells reduces or enhances ISO-induced CM hypertrophic growth, respectively (Fig. 2h, i). To confirm the in-trans effects of miR-574 in cardioprotection, we injected miR-574-5p/3p mimics into miR-574^−/−^ mice and could significantly rescue the cardiac disease phenotype under ISO treatment including reduced CM hypertrophy, fibrosis (accompanied by reduced hypertrophy and fibrosis marker gene expression), and myocyte death (Supplementary Fig. 9). More detailed functional characterizations of both miRNAs in the two cardiac cell types need to be further performed in the future.

miR-574^−/−^ mice are normal in the absence of stress for two possible reasons: 1) redundant miRNAs that share common target seed sequences compensate for the loss of function of genetically deleted miR-574, or 2) compensatory adaptive responses in the miR-574 knockout mouse offset the loss-of-function effect. The latter may be a major reason for the absence of baseline phenotypes in miR-574^−/−^ mice, as multiple MEGs are reduced at the transcriptional level. These transcriptional alterations may restore the balance after FAM210A induction and consequent increased expression of MEG proteins, preventing dramatic phenotypical changes. We identified bona fide miR-574-5p/3p targets by performing RNA-Seq of the total RNA from the hearts of WT and miR-574^−/−^ mice at baseline. We found that 34 genes are significantly upregulated at the mRNA level. Among these, 8 genes contain the putative miR-574-5p target seed sequence, while one single gene contains the putative miR-574-3p target seed sequence. Although FAM210A is identified as the only common target gene of both miR-574-5p and miR-574-3p in our studies, the function and contribution of other putative targets of miR-574-5p/3p warrant further investigation.

RNA-binding proteins (RBPs) and miRNAs play key roles in posttranscriptional regulation of mRNAs such as mRNA stability and translatability^50^. Our recent report suggests that RBPs can bind to miRNAs as a “sponge” in a reversed direction and regulate their function. We have shown that an RBP heterogeneous ribonucleoprotein L (hnRNP L) forms a complex with miR-574-3p via binding a CA-rich element and inactivates Ago2 loading and the miRNA activity in monocytes^48^. Intriguingly, Ago2 CLIP-Seq barely detected any miR-574-5p and miR-574-3p binding to Ago2 in advanced cardiomyopathy patients, despite its induced expression in both ischemic and non-ischemic HF patients, as indicated by comparative miRNA-Seq analyses of myocardial miRNAs^51, 52^. This finding implies that both strands of miR-574 may be eliminated from the RISC by an unknown mechanism. TDP-43 is the top four RNA-binding proteins (RBPs) identified in HL-1 CM cells by mRNA Interactome Capture analyses.^53^ TDP-43 binds to GU-rich mature miR-574-5p *in vitro*^54, 55^, and inactivates miR-1/206 through direct binding in the C2C12 skeletal muscle cell line.^56^ We detected an interaction between TDP-43 and miR-574-5p in the late stage (but not in the early stage) of cardiac hypertrophy by the ribonucleoprotein immunoprecipitation (RIP) coupled with RT-qPCR, consistent with the exclusion from Ago2 binding at late-stage cardiac remodeling (Supplementary Fig. 10a, b). As a summary, we conclude that the cardiac stress-induced miR-574-5p protects CMs from pathological cardiac hypertrophy at the early stage of cardiac remodeling but may undergo inactivation by TDP-43 in the late stage. Similarly, miR-574-3p may be inactivated by hnRNP L in a late stage of HF as well. Alternatively, hnRNP L-miR-574-3p complex may have a distinct Ago2-independent function. Interestingly, 45 human single nucleotide polymorphisms (SNPs) are present in the miR-574 gene (Supplementary Fig. 1d, Ensembl genome browser). Three mutations delete the GU dinucleotide within the seed region of miR-574-5p, and four point mutations change the CA-rich element CACA into UACA, CAUA or CACG in the 3′-end of miR-574-3p. Some of these SNPs reach high frequency (MAF=0.001) and may alter the *in vivo* function of miR-574 and increase the risk for triggering cardiac disorders.

A genome-wide meta-analysis shows that the genetic variations near FAM210A are associated with reduced lean mass and increased risk of bone fracture^38, 57^. A prior study of FAM210A explored the transcriptional profile in primary muscle cells isolated from skeletal muscle-specific Fam210a conditional KO mouse and found that multiple muscle differentiation-related genes were decreased^38^. However, the molecular mechanism by which FAM210A regulates muscle functions is still unknown. In this study, we discover that FAM210A regulates MEG protein expression based on polysome profiling and AHA labeling assays. The mechanism of the regulation is currently under investigation based on several clues from our data and literature. FAM210A, the mitochondrial elongation factor EF-Tu, and ATAD3A form a complex. ATAD3A is an essential protein for mitochondrial protein synthesis^41^. Mass spectrometry screens showed that both FAM210A and ATAD3A are associated with the mitoribosome^41^. FAM210A interacts with EF-Tu and may promote translation elongation by promoting the delivery of aminoacyl-tRNAs^58^ (Supplementary Fig. 7f). We speculate that ATAD3A may hydrolyze ATP and provide energy for FAM210A to recruit aminoacyl-tRNA-bound EF-Tu and dissociate EF-Tu when aminoacyl-tRNA enters the A-site of mitoribosome.

Our study suggests that FAM210A may contribute to coordinating the expression of MEG proteins and NEMG proteins in CMs (Supplementary Fig. 7f). However, under cardiac stress and tissue injury, FAM210A exaggerates the pathological remodeling response by over-activating mitochondrial protein expression and altering cytoplasmic-mitochondrial ETC component protein balance. These responses may promote HF progression. FAM210A is ubiquitously expressed in multiple cell types (CMs, CFs, skeletal muscle cells, HEK293T, etc.) and organs (heart, skeletal muscle, brain, etc.)^38^, implying a conserved and generalized role for this protein. Similar to FAM210A, miR-574 is ubiquitously expressed and more enriched in heart, brain, and skeletal muscle (Supplementary Fig. 1a). Thus, the miR-574-mediated regulation of FAM210A may also play important roles in maintaining mitochondrial protein homeostasis and mitochondrial activity in other organs and related diseases. Our studies provide novel insights into the function and regulatory mechanisms of FAM210A in the modulation of MEG protein expression and mitochondrial activity. In the future, we will explore the cardiac function of FAM210A using cardiomyocyte-specific knockout and transgenic mouse models.

Translation of mitochondrial-encoded genes needs to be tightly regulated to synchronize with that of nuclear-encoded mitochondrial genes in order to maintain mitochondrial homeostasis^7^. To date, translational control mechanisms of MEGs are largely unknown. Previous studies revealed that miR-1 and miR-21 enter mitochondria and directly bind to several MEG mRNAs to activate their translation in skeletal muscle cells^31^ and cardiac myocytes^32^, respectively. A recent integrated omics analysis of proteomic changes during cardiac remodeling has shown that the mitochondrial translation elongation pathway is involved in cardiac hypertrophy^59^. However, the underlying mechanism is unknown. In this study, we have uncovered a novel mechanism by which the miR-574-FAM210A axis regulates MEG protein expression in cardiac mitochondria and potentially in other tissues or cell types (Fig. 7i, Supplementary Fig. 7f).

Proteomic studies show that the steady-state level of multiple MEG proteins in the ETC complex was increased in the early stage of cardiac hypertrophy in multiple HF models^60^. Data-mining and re-analyses of TRAP-Seq results lead us to discover that after TAC, nuclear-encoded ETC genes were mostly upregulated at the translational level in ACMs (Supplementary Fig. 7g)^8^. Using a human cardiomyocyte cell culture system, we show that miR-574 regulates MEG protein expression by targeting FAM210A after manipulating the ratio of miR-574 and FAM210A in an acute phase (Fig. 7). These results imply a coordinated expression of MEG proteins by FAM210A in response to elevated NEMG protein expression to maintain the stoichiometric balance. miR-574-5p and miR-574-3p may function as a potential checkpoint molecule to prevent excessive MEG protein expression (overcompensation), maintain mitochondrial homeostasis, and reduce CM hypertrophy. To determine the change of mitochondrial protein expression *in vivo*, we also measured MEG and NEMG protein expression in TAC and Sham hearts from WT versus miR-574^−/−^ mice. We showed that expression of FAM210A protein is higher in miR-574^−/−^ hearts compared to WT hearts 1-week post-TAC surgery (Supplementary Fig. 8h). However, MEG proteins (such as ND1 and COX2) are not significantly increased in miR-574^−/−^ hearts compared to WT hearts under TAC surgery. These data suggest that intracellular communication and autonomous secondary effects such as the mitochondrial protein degradation pathway might affect protein expression of MEG beyond translational regulation and lead to re-balance between MEG and NEMG protein expression in a more chronic phase.

miR-574 is induced by hypertrophic stress as an anti-hypertrophy molecule in CMs; it targets *Fam210a* mRNA, among other transcripts. Thus, we have shown that two strands of a specific miRNA gene fine-tune the expression level of FAM210A, which may directly or indirectly regulate mitochondrial encoded protein expression. These findings offer novel insights into the development of therapeutic approaches for the treatment of heart failure. We propose that normalizing mitochondrial translation (removing an excessive amount of translation activity) using miR-574-5p and miR-574-3p could be used to antagonize cardiac pathological remodeling in the treatment of heart disease.

## Methods

### Human specimens

All human samples of frozen cardiac tissues (n=23), including 17 samples from explanted failing hearts and 6 samples from donor non-failing hearts, were acquired from the Cleveland Clinic. This study was approved by the Material Transfer Agreement between the URMC and the Cleveland Clinic. Total RNA was extracted from human heart tissues using TRIzol reagent (ThermoFisher Scientific) following instructions in the manual. For the mRNA detection, 1 μg of total RNA was used as a template for reverse transcription assay using the iScript cDNA Synthesis Kit (Bio-Rad); cDNA was used for detecting FAM210A and GAPDH expression. For the expression of miRNA, the total RNA (containing small RNAs) was subjected to miRNA reverse transcription using the miScript II RT Kit (Qiagen), following the manual, and the RT product was used for detection of miR-574-5p/3p and control RNA Snord68 using the miScript Primer Assay (Qiagen).

### Mice

Three Sanger MirKO ES cell lines for Mir574 (produced by Wellcome Trust Sanger Institute)^35^ were purchased from the mutant mouse regional resource center (MMRRC). miR-574^−/−^ global knockout chimera founder mice were generated in the Case Western Reserve University Transgenic Core Facility. We generated the miR-574 targeted male chimera mouse in the C57BL/6J background and performed germline transmission and backcrossed the mice to C57BL6J mice for more than 10 generations. A 182-bp DNA region was deleted, including the microRNA574 gene (96-bp for pre-miR-574 sequence) in the intron 1 of the host gene Fam114a1. Puromycin resistance cassette was removed by breeding with C57BL/6-Tg(Zp3-cre)93Knw/J mouse, which expresses Cre recombinase in oocytes, resulting in a null allele. For experiments with miR-574^−/−^ mice, control mice of the same age and gender from littermates or sibling mating were used. All animal procedures were performed in accordance with the National Institutes of Health (NIH) and the University of Rochester Institutional guidelines.

### Isoproterenol (ISO) injection and infusion model

Experimental mice are siblings generated from intercrosses of miR-574^+/–^ mice. Age and background matched WT and miR-574^−/−^ female mice at 10-12 weeks of age were subjected to a vehicle (saline) or ISO treatment. The procedure for ISO injection: ISO or vehicle saline were administered to WT and miR-574^−/−^ mice daily on weekdays for 4 weeks, using subcutaneous injection (30 mg/Kg/day). Excised mouse hearts were washed with saline to remove the blood, fixed in 10% formalin, and used for histological and immunoblotting analyses. The procedure for mini-osmotic pump implantation: Mouse is anesthetized using 2.0% isoflurane and placed on a heated surgical board. A side/upper back area skin incision is made, and the mini-osmotic pump is inserted subcutaneously set to deliver ISO or vehicle. The incision is then closed with 6-0 coated vicryl in a subcuticular manner, and the animals are allowed to recover. The pumps will not be removed and will remain for a period of 4 weeks, and the animals will be euthanized. The sutures will be removed after 2 weeks since the pumps are transplanted.

### Transverse aortic constriction (TAC) surgical model

Experimental mice are siblings generated from intercrosses of miR-574^+/–^ mice. Age and background matched WT and miR-574^−/−^ male mice at 10-12 weeks of age were subjected to Sham or TAC surgery.

The mouse is anesthetized using 2.0% isoflurane, placed on a surgical board on a heating pad (half-inch plexiglass between the animal and the heating pad), and given buprenorphine SQ. A midline cervical incision is made to expose the trachea for visualizing oral intubation using a 22-gauge (PE90) plastic catheter. The catheter is connected to a volume-cycled ventilator supplying supplemental oxygen with a tidal volume of 225-250 μl and a respiratory rate of 120-130 strokes/min. Surgical plane anesthesia is subsequently maintained with 1-1.5% isoflurane. A left thoracotomy is performed: Skin is incised and chest cavity opened at the level of the 2^nd^ intercostal space. Transverse section of the aorta is isolated. Transverse aortic constriction is created by placing a (6-0 silk) ligature securely around the trans-aorta and a 27-gauge needle, causing complete occlusion of the aorta. The needle is removed, restoring a lumen with severe stenosis. Lungs are reinflated and the chest is closed using Vicryl 6-0 suture. Muscle and skin are sutured using a Vicryl 6-0 suture in a running subcuticular pattern. Once the mouse is breathing on its own, it is removed from the ventilator and allowed to recover in a clean cage on a heated pad.

### Cell culture

AC16 adult human ventricle cardiomyocyte cells, HL-1 mouse atrial myocyte cells, and HEK293T cells were used to address questions at the cellular and molecular levels. Both cells were cultured following the manual (Sigma, Cat. No. SCC109 for AC16, SCC065 for HL-1). AC16 cells were cultured in DMEM/F12 containing 2 mM L-Glutamine, 12.5% FBS, and 1x Penicillin/Streptomycin Solution. For the myocyte hypertrophy experiment, AC16 cells were treated with 10 μM ISO for 24 hrs. HL-1 cells were cultured in Claycomb medium containing 2 mM L-Glutamine, 10% FBS, 1x Penicillin/Streptomycin Solution, and 0.1 mM Norepinephrine on a pre-coated plate using 0.25 mM fibronectin in 0.02% gelatin. HEK293T cells were cultured in DMEM containing 2 mM L-Glutamine, 10% FBS, 1x Penicillin/Streptomycin Solution.

### Luciferase activity assay

Reporter vectors were generated by inserting the Fam210a 3’UTR fragments containing wild-type or mutated seed sequence into the miRNA reporter vector pmirGlo (Promega). All the primers used to generate the reporter vector were listed in supplemental information. The wild-type 3’UTR fragment was amplified and ligated to pmirGlo. Then, 100 ng of wild-type or mutant reporter vectors were co-transfected with 20 nM miRNA (miR-574-5p or miR-574-3p) into HEK293T cells cultured in 24-well plate using lipofectamine 3000 (ThermoFisher Scientific) following the manufacturer’s instruction. Cells were collected at 36 hrs after transfection, and firefly and renilla luciferase activity were detected by the Dual-Luciferase Reporter Assay System (Promega).

### AHA labeling assay

Click-iT reagents (ThermoFisher Scientific) were used to label newly synthesized polypeptides in mitochondria as previously described^31^. Briefly, HEK293T cells were infected with FAM210A shRNA or overexpression lentivirus and cultured for 24 hrs. The cells were washed 3 times with pre-warmed PBS and incubated in pre-warmed methionine-free DMEM for 45 mins followed by the addition of 20 μM Emetine (Sigma, E2357) for 15 mins to block the cytosolic protein synthesis. Then the medium was replaced with methionine-free DMEM containing 50 μM methionine analog AHA and 20 μM Emetine, and cells were incubated for 4 hrs. The AHA incorporated newly synthesized proteins in mitochondria were labeled with TAMRA by Click-iT reaction following the instruction. Labeled proteins were separated by 4-15% gradient SDS-PAGE, and analyzed by the ChemiDoc MP system (Bio-Rad). The protein gel was also subject to immunoblot for TOM20 as a loading control.

### Analysis of Cytosolic and Mitochondrial Translational Activity by Polysome Profiling

Polysome profiling for the nuclear-encoded mitochondrial genes (cytosolic translation) was performed as previously described^48^. Cycloheximide (CHX, 100 μg/ml) was added to the cells for 15 mins before lysis to freeze ribosomes on mRNAs in the elongation phase. Around 10^7^ cells were lysed in TMK lysis buffer (10 mM Tris-HCl pH 7.4, 100 mM KCl, 5 mM MgCl2, 1% Triton X-100, 0.5% Deoxycholate, 2 mM DTT) containing 100 μg/ml CHX, 4 U/ml RNase inhibitor (NEB) and proteinase inhibitor cocktail (Roche) on ice for 20 mins. Equal amounts of A260 absorbance from each sample were loaded onto a 10-50% sucrose gradient solution and centrifuged at 29,000 rpm for 4 hrs; 22 translation fractions were collected from each sample by Density Gradient Fractionation System (BRANDEL). Based on the UV absorbance curve, the 22 fractions were pooled into 8 samples, including free mRNP, 40S small ribosome subunit, 60S large ribosome subunit, 80S monosome, light polysomes (di-ribosome, tri-ribosome, etc.), and heavy polysomes (>5 ribosomes). Total RNA was extracted from the same volume of each pooled fraction with Trizol LS (ThermoFisher Scientific), and Renilla luciferase mRNA from *in vitro* transcription was used as RNA spike-in and loading control for RT-qPCR. The mitochondrial polysome profiling was performed as previously described^61^. The mitochondria were isolated from cells and lysed in TMK lysis buffer. Then the samples were loaded onto a 10-30% sucrose gradient solution and centrifuged following the same procedure as with cytosolic polysome profiling. 12 pooled fractions were collected from mitochondrial polysome profiling for further analysis of the translation of mitochondrial-encoded genes by RT-qPCR. 12S and 16S small and large rRNAs were used as quality controls for polysome profiling. The association of mitochondrial-encoded ETC gene mRNAs was analyzed by RT-qPCR.

#### Interactome capture by Immunoprecipitation-Mass spectrometry (IP-MS)

1. Immunoprecipitation: The total mitochondria were isolated from the whole heart of wild-type mice following the manual using the Mitochondria Isolation Kit for Tissue (Pierce). The mitochondria pellets were re-suspended in NP-40 lysis buffer (50 mM HEPES, pH 7.5, 150 mM KCl, 0.5% NP-40, 2 mM EDTA, 1 mM NaF, 0.5 mM DTT with proteinase inhibitor cocktail), and incubated on ice for 15 mins. The mitochondrial lysate was centrifuged at 12,000 rpm for 10 mins. The suspension was equally divided into two parts and incubated with 1 μg rabbit pre-immune IgG or anti-FAM210A rabbit polyclonal antibody at 4°C with rotation overnight, respectively. The protein-antibody complex was pulled down by incubation with Dynabeads Protein G (ThermoFisher Scientific) for 4 hrs and eluted using 1x SDS loading buffer. For the mass spectrometry assay, the elution was loaded onto 4-12% gradient SDS-PAGE gel and run for 10 mins. The whole lane was subjected to mass spectrometry analysis by Mass Spectrometry Resource Lab of the University of Rochester Medical Center. The interaction of FAM210A and EF-Tu or ATAD3A was confirmed by Western blot following IP using indicated antibodies.
2. Sample Preparation: For mass spectrometry experiments, samples were run into a 4-12% SDS-PAGE gel for a short time to remove contaminants and create a ∼10 mm length region, allowing the total protein to be evaluated in a single gel digest. After staining with SimplyBlue SafeStain (Invitrogen), these regions were excised, cut into 1 mm cubes, de-stained, then reduced and alkylated with DTT and IAA, respectively (Sigma). Gel pieces were dehydrated with acetonitrile. Aliquots of trypsin (Promega) were reconstituted to 10 ng/μl in 50 mM ammonium bicarbonate and added so that the solution was just covering the dehydrated gel pieces. After 0.5 hr at room temperature (RT), additional ammonium bicarbonate was added until the gel pieces were completely submerged and placed at 37°C overnight. Peptides were extracted the next day by adding 0.1% TFA, 50% acetonitrile, then dried down in a CentriVap concentrator (Labconco). Peptides were desalted with homemade C18 spin columns, dried again, and reconstituted in 0.1% TFA.
3. FAM210A-IP LC-MS/MS: Peptides were injected onto a homemade 30 cm C18 column with 1.8 μm beads (Sepax), with an Easy nLC-1000 HPLC (ThermoFisher Scientific), connected to a Q Exactive Plus mass spectrometer (ThermoFisher Scientific). Solvent A was 0.1% formic acid in the water, while solvent B was 0.1% formic acid in acetonitrile. Ions were introduced to the mass spectrometer using a Nanospray Flex source operating at 2 kV. The gradient began at 3% B and held for 2 mins, increased to 30% B over 41 mins, increased to 70% over 3 mins and held for 4 mins, then returned to 3% B in 2 mins and re-equilibrated for 8 mins, for a total run time of 60 mins. The Q Exactive Plus was operated in a data-dependent mode, with a full MS1 scan followed by 10 data-dependent MS2 scans. The full scan was done over a range of 400-1400 m/z, with a resolution of 70,000 at m/z of 200, an AGC target of 1e6, and a maximum injection time of 50 ms. Ions with a charge state between 2-5 were picked for fragmentation. The MS2 scans were performed at 17,500 resolution, with an AGC target of 5e4 and a maximum injection time of 120 ms. The isolation width was 1.5 m/z, with an offset of 0.3 m/z, and a normalized collision energy of 27. After fragmentation, ions were put on an exclusion list for 15 seconds to allow the mass spectrometer to fragment lower abundant peptides.
4. Data Analysis: Raw data from MS experiments were searched using the SEQUEST search engine within the Proteome Discoverer software platform, version 2.2 (ThermoFisher Scientific), using the SwissProt human database. Trypsin was selected as the enzyme allowing up to 2 missed cleavages, with an MS1 mass tolerance of 10 ppm. Samples run on the Q Exactive Plus used an MS2 mass tolerance of 25 mmu. Carbamidomethyl was set as a fixed modification, while oxidation of methionine was set as a variable modification. The Minora node was used to determine relative protein abundance between samples using the default settings. Percolator was used as the FDR calculator, filtering out peptides which had a q-value greater than 0.01.

### Immunofluorescence and confocal microscopy

Immunostaining of cells grown on coverglass or chambered slides: HEK293T, AC16 cells, primary CMs were grown on the coverslips, and incubated with 100 nM MitoTracker Red CMXRos (ThermoFisher Scientific) for 30 mins at 37°C before being fixed for 10 mins with 4% paraformaldehyde in PBS. Cells were washed by PBS for 3x 5 mins and permeabilized with ice-cold 0.5% Triton X-100 in PBS for 5 mins. After blocking with 1% BSA in PBS, the coverslips were incubated with indicated primary antibodies (anti-FAM210A 1:1000; anti-Flag 1:2000) in blocking solution (2% BSA in PBS) for 1 hr at RT and washed with PBS for 3x 5 mins. Then, the coverslips were incubated with the Alex Fluor-488 conjugated secondary antibodies (ThermoFisher Scientific, 1:1000) in PBS and washed with PBS for 3x 5 mins. Coverslips were air dried and placed on slides with antifade mounting medium (containing DAPI). The slides were imaged using an Olympus FV1000 confocal microscope.

### Wheat germ agglutinin (WGA) and phalloidin staining

WGA (5 mg) was dissolved in 5 ml of PBS (pH 7.4). We performed deparafinization by following steps: **a.** Xylene (100%) for 2x 5 mins; **b.** Ethanol (100%) for 2x 5 mins; **c.** Ethanol (95%) for 1x 5 mins; **d.** ddH2O for 2x 5 mins. The slides were kept in a pressure cooker for 10 mins along with citrate buffer (10 mM, pH 6.0) for antigen retrieval. We quenched the slides with 0.1 M glycine in phosphate buffer (pH 7.4) for 1 hr at RT. Circles were made with Dako pen, and slides were blocked with goat normal serum for 30 mins. 10 μg/ml of WGA-Alexa Fluor 488 (Sigma Aldrich) was applied to the slides for incubation for 1 hr at RT. Slides were rinsed in PBST wash buffer 3x 5 mins followed by PBS for 5 mins. We placed a coverslip on the slides with an antifade solution (containing DAPI) for imaging. Five different cross-sectional areas were selected, and the cell size of at least 500 CM cells was measured per area. For primary murine CM cell culture, Alexa Fluor^TM^ 594 Phalloidin (ThermoFisher Scientific, Cat. No. A12381) was used to measure the cell size following the instruction from the manual. Primary CM cells were treated with 10 mM ISO for 24 hrs for measuring cell size using Phalloidin and for 48 hrs for the TUNEL assay, respectively. Cultured CMs were fixed using 4% paraformaldehyde in PBS for 10 mins, washed in PBS, and permeabilized in 0.2% Triton X-100 for 10 mins. Cells were blocked in 2% BSA/PBS for 1 hr and stained with Alexa Fluor^TM^ 594 Phalloidin in 1:1000 dilution for 30 mins at RT. The stained cells were gently washed with PBS for 3x 5 mins, and the slides were mounted using a mounting medium with DAPI.

### Picrosirius red staining

Paraffin-embedded tissue sections were deparaffinized and incubated in a picrosirius red solution (Abcam, Cat. No. ab150681) at RT for 1 hr. Then, slides were subjected to 2 washes of 1% acetic acid and 100% of ethyl alcohol and mounted in a resinous medium. Images were captured using the PrimeHisto XE Histology Slide Scanner (Carolina), for the analysis six images were selected from each group. Total collagen content was determined for the whole heart images using an Image J software.

### Terminal deoxynucleotidyl transferase dUTP nick end labeling (TUNEL) staining

Primary adult CMs were plated and grown on glass slides with chambers. The cells were washed with PBS twice and fixed with 4% paraformaldehyde for 20 mins. Cells were permeabilized with 0.5% of Triton X-100 for 5 mins, and incubated with TUNEL reaction mixture (In Situ Cell Death Detection Kit; Sigma, 11684795910) for 1 hr at 37°C in the dark. Finally, cells were washed with PBS for 3x 5 mins, air dried, and mounted with DAPI-containing antifade medium. Images were captured using a BX51 microscope (Olympus).

### Determination of cellular ATP content

The cellular ATP content was determined using a Molecular Probes ATP Determination Kit (ThermoFisher Scientific, A22066) according to the manufacturer’s instructions. Briefly, cells were incubated in 1% (w/v) of Trichloroacetic acid and centrifuged at 10,000 rpm for 5 mins at 4°C. The supernatant was collected and neutralized by Tris-HCl (0.1 M, pH 9.0), then mixed with the bioluminescent reagent. After incubation, bioluminescent signals were read using the HTX microplate reader (BioTek Instruments). ATP concentrations were determined in all samples based on a standard curve and normalized by the total protein mass.

### Measurement of mitochondrial membrane potentiality

The mitochondrial membrane potentiality was measured using a TMRE (Tetramethylrhodamine ethyl ester, ThermoFisher Scientific, Cat. No. T669) according to the manufacturer’s protocol. Isolated adult mouse CMs were loaded with TMRE at 25 nM concentration for 30 mins at RT. After incubation, cells were washed with 1x PBS for 3 times, and cell culture dish filled with the live cell imaging solution and fluorescence images were captured using a laser scanning confocal microscope (Olympus).

### Dihydroethidium (DHE) staining

DHE staining was performed in both isolated live murine cardiomyocyte cells and frozen heart tissue sections. Primary adult CMs were isolated from WT and miR-574^−/−^ mice and cultured using the standard procedure. Also, heart sections were prepared by the frozen section procedure from WT and miR-574^−/−^ mice. In situ superoxide radical’s production in live CMs or frozen sections was measured with the oxidative fluorescent dye called Dihydroethidium (DHE, Thermofisher Scientific). The equal volume of cells was plated and then treated with vehicle (0.5% DMSO) and ISO for 24 hrs. The cell sections were incubated with 10 μM DHE for detection of intracellular reactive oxygen species (ROS) for 20 mins at 37°C in the dark and washed with PBS for 3x 15 mins. Sections were air-dried and mounted with a coverslip. Images were captured using an Upright digital immunofluorescence microscope (BX51, Olympus). Four images were taken from each section. Fluorescence signal intensities were quantified with Image J software.

### Statistical Analysis

All quantitative data were presented as mean ± SEM and analyzed using Prism 7 software (GraphPad). For a comparison between 2 groups, a Student *t* test was performed. For multiple comparisons among ≥3 groups, 1-way ANOVA was performed. Statistical significance was assumed at a value of *P*≤0.05. Further details and a complete collection of methods are available in the Online Data Supplement.

## Supplementary Figure Legends

**Supplementary Fig. 1.**
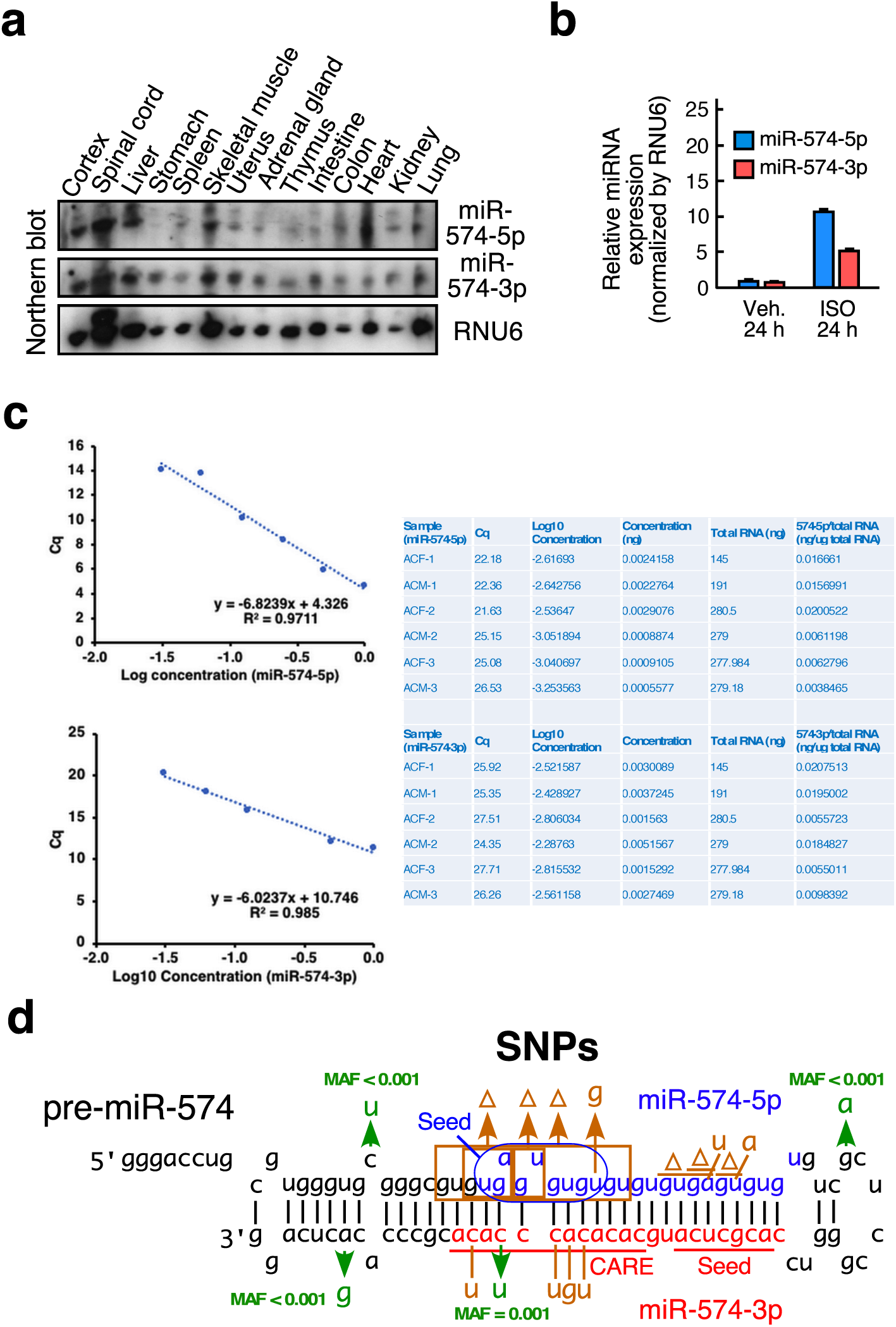
Baseline expression of miR-574-5p/3p in murine organs and ACMs, and the structure of miR-574 precursor. **a** Northern blot detection of miR-574-5p/3p expression in various murine organs. **b** ISO induces miR-574-5p and miR-574-3p expression in isolated murine ACMs. **c** Quantitative measurement of absolute copy number of miR-574-5p and miR-574-3p in freshly isolated CM and CF cells. Left panel: Standard curve of synthesized miR-574-5p/3p by RT-qPCR. Right panel: RT-qPCR for miR-574-5p/3p from total RNA extracted from murine CM and CF cells immediately after isolation. **d** The schematic of the secondary structure of miR-574 precursor containing guide strand miR-574-5p and passenger strand miR-574-3p. Out of 45 human SNPs uncovered in the miR-574 precursor, multiple SNPs with high frequency or located in mature miR-574-5p/3p sequences were highlighted.

**Supplementary Fig. 2.**
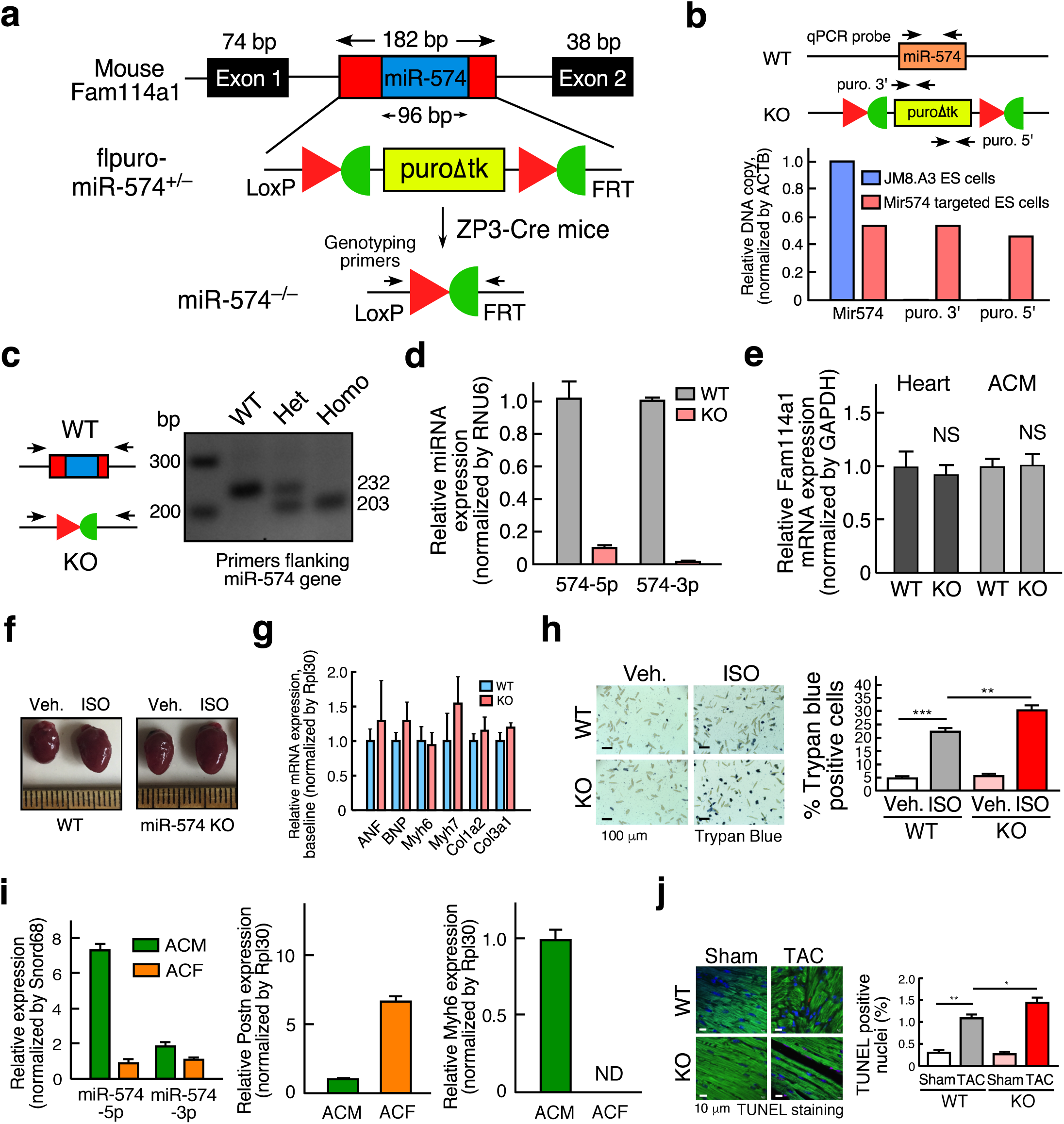
Generation of miR-574 global knockout mice and phenotypic characterization. **a** The schematic of the generation of miR-574^−/−^ mice. **b** Characterization of Mir574 targeted ES cells for the construction of KO mice. **c** Genotyping of miR-574^−/−^ mice with removed puromycin resistance cassette. **d** miR-574-5p/3p expression in WT and miR-574^−/−^ mice, n=3. **e** The expression of host gene *Fam114a1* mRNA in the heart and CMs of WT and miR-574^−/−^ mice. NS, non-significant. **f** Pictures of hearts from WT and miR-574^−/−^ mice with the vehicle and ISO treatment. **g** RT-qPCR of fetal cardiac genes in WT and miR-574^−/−^ mice at baseline. **h** Trypan blue staining of primary ACMs from WT and miR-574^−/−^ mice under ISO versus vehicle treatment. **i** Left panel: Expression of miR-574-5p/3p in primary murine ACMs and ACFs. Middle and right panels: RT-qPCR of CM and CF marker genes for quality control of cardiac cell isolation. **j** TUNEL assay for heart tissue sections from WT and miR-574^−/−^ mice under Sham versus TAC surgery. Data were presented as mean ± SEM. *: p≤0.05, **: p≤0.01, ***: p≤0.001 by Student *t* test or one-way ANOVA.

**Supplementary Fig. 3.**
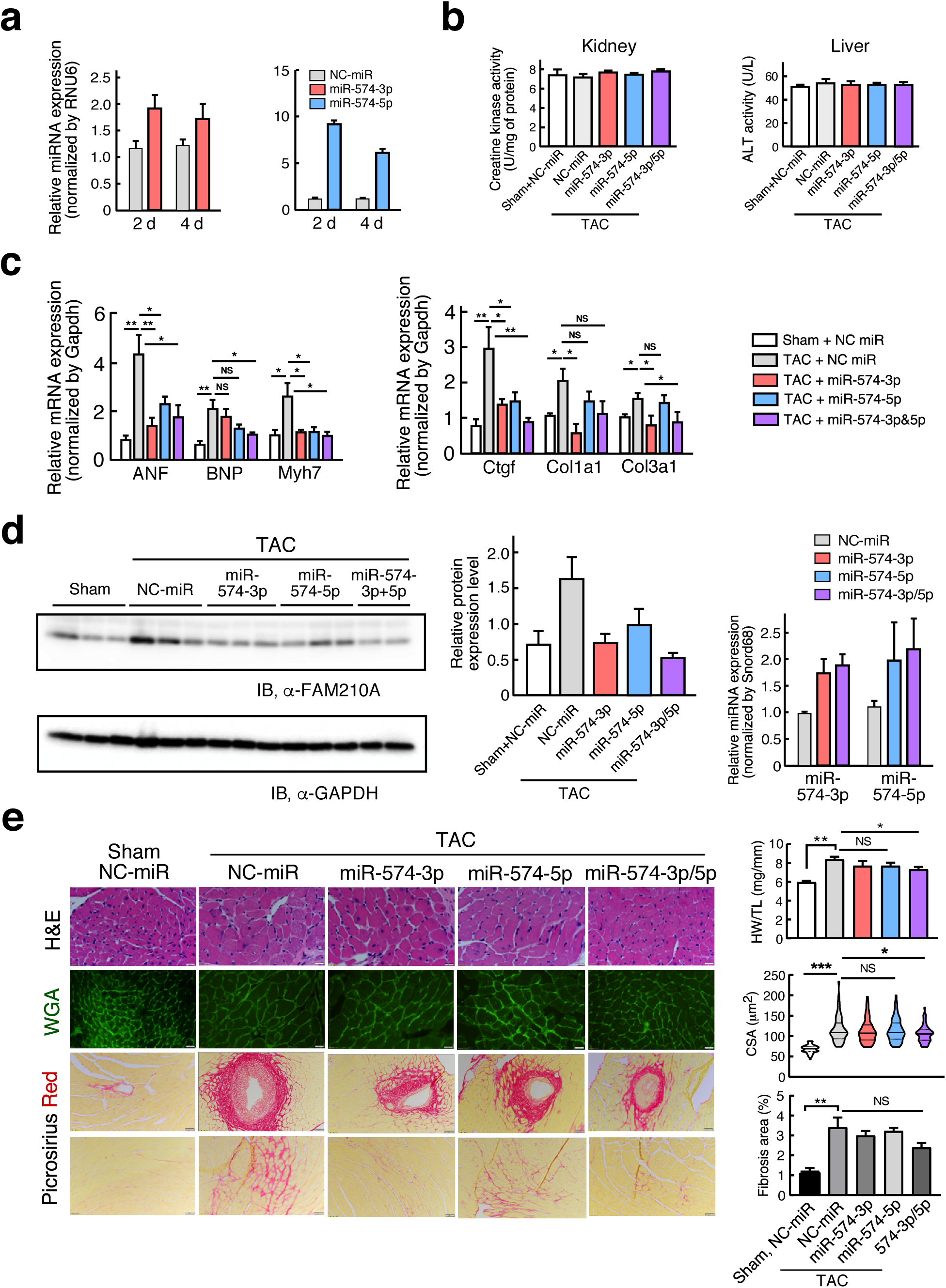
Phenotypic characterization of the therapeutic model using miR-574-5p/3p mimic injection in mice subject to TAC surgery. **a** RT-qPCR of miR-574-5p and miR-574-3p after injections of the miRNA mimics for 2 or 4 days. **b** Creatine kinase test and alanine transaminase (ALT) assay in kidneys and livers from miRNA mimic injected mice. **c** RT-qPCR of hypertrophy and fibrosis marker genes in the hearts of therapeutic mouse models. **d** Western blotting of FAM210A in the hearts of therapeutic mouse models. Expression of miR-574-5p and miR-574-3p was measured by RT-qPCR at the end-point. **e** Low dose (1 mg/Kg BW) miR-574-5p/3p injection does not show significant cardioprotection. *: p≤0.05, **: p≤0.01 by one-way ANOVA.

**Supplementary Fig. 4.**
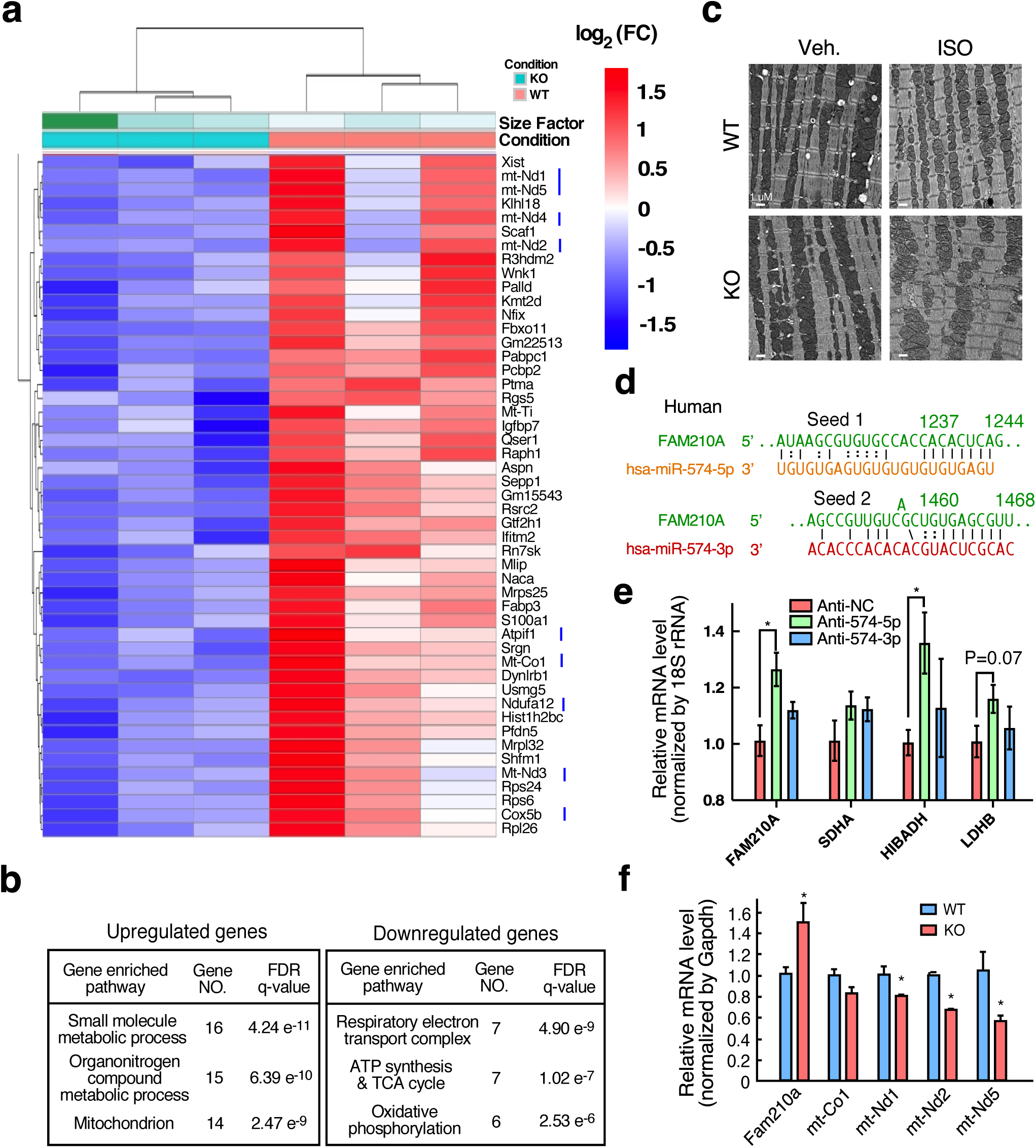
Downregulated genes in miR-574 KO versus WT mouse heart and validation of miR-574 target genes. **a** Heatmap of significantly downregulated genes in miR-574^−/−^ mice at baseline analyzed by RNA-Seq. P60 male mice, n=3 per group, padj<0.05. **b** Gene Ontology analysis of enriched pathways of dysregulated genes in RNA-Seq. The top three pathways are listed with enriched gene sets. **c** Electron microscopic imaging of mitochondria in CMs from hearts of WT and miR-574^−/−^ mice. **d** Seed sequence sites of miR-574-5p and miR-574-3p in human *FAM210A* mRNA. **e** Validation of miR-574-5p target genes using RT-qPCR analysis of mRNA expression in AC16 cells transfected with control anti-miR or anti-miR-574-5p/3p inhibitors. The statistical calculation is performed in comparison with the control anti-miR transfection sample. **f** RT-qPCR validation of multiple dysregulated genes in hearts from miR-574^−/−^ versus WT mice. *: p≤0.05, **: p≤0.01 by Student *t* test or one-way ANOVA.

**Supplementary Fig. 5.**
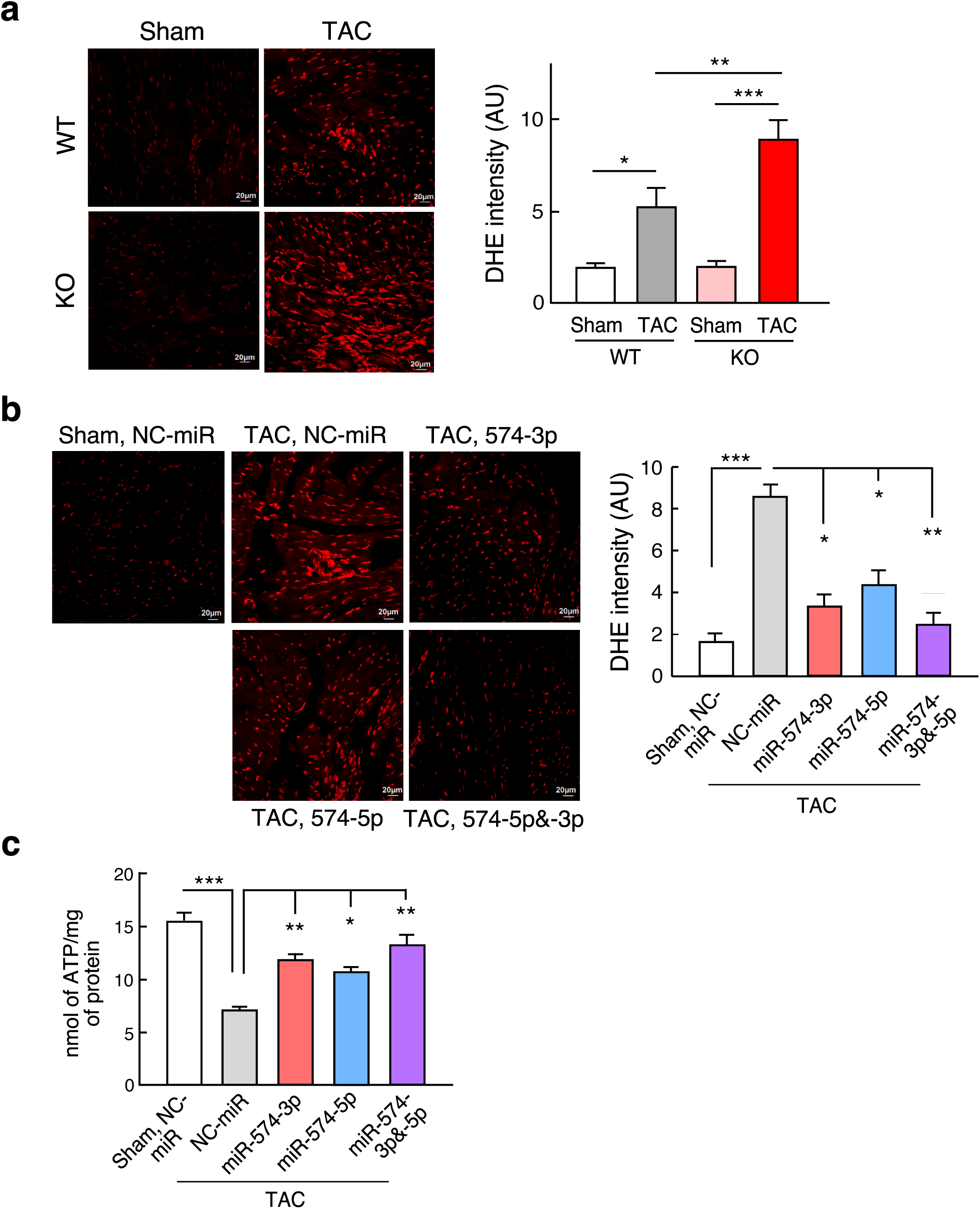
Assessment of mitochondrial functions in genetic, disease, and therapeutic mouse models. **a** DHE staining of frozen sections of hearts from WT and miR-574^−/−^ mice under Sham and TAC operations (4 weeks post-surgery). **b** DHE staining of frozen sections of hearts from WT mice under treatment with miRNA mimics in TAC-induced HF mouse models. NC-miR: negative control miRNA mimics. Sham operation was used as a control for TAC (6 weeks post-surgery). **c** ATP production in heart lysates from WT mice under treatment with miRNA mimics in TAC-induced HF mouse models (6 weeks post-surgery). *: p≤0.05, **: p≤0.01, ***: p≤0.001, by one-way ANOVA.

**Supplementary Fig. 6.**
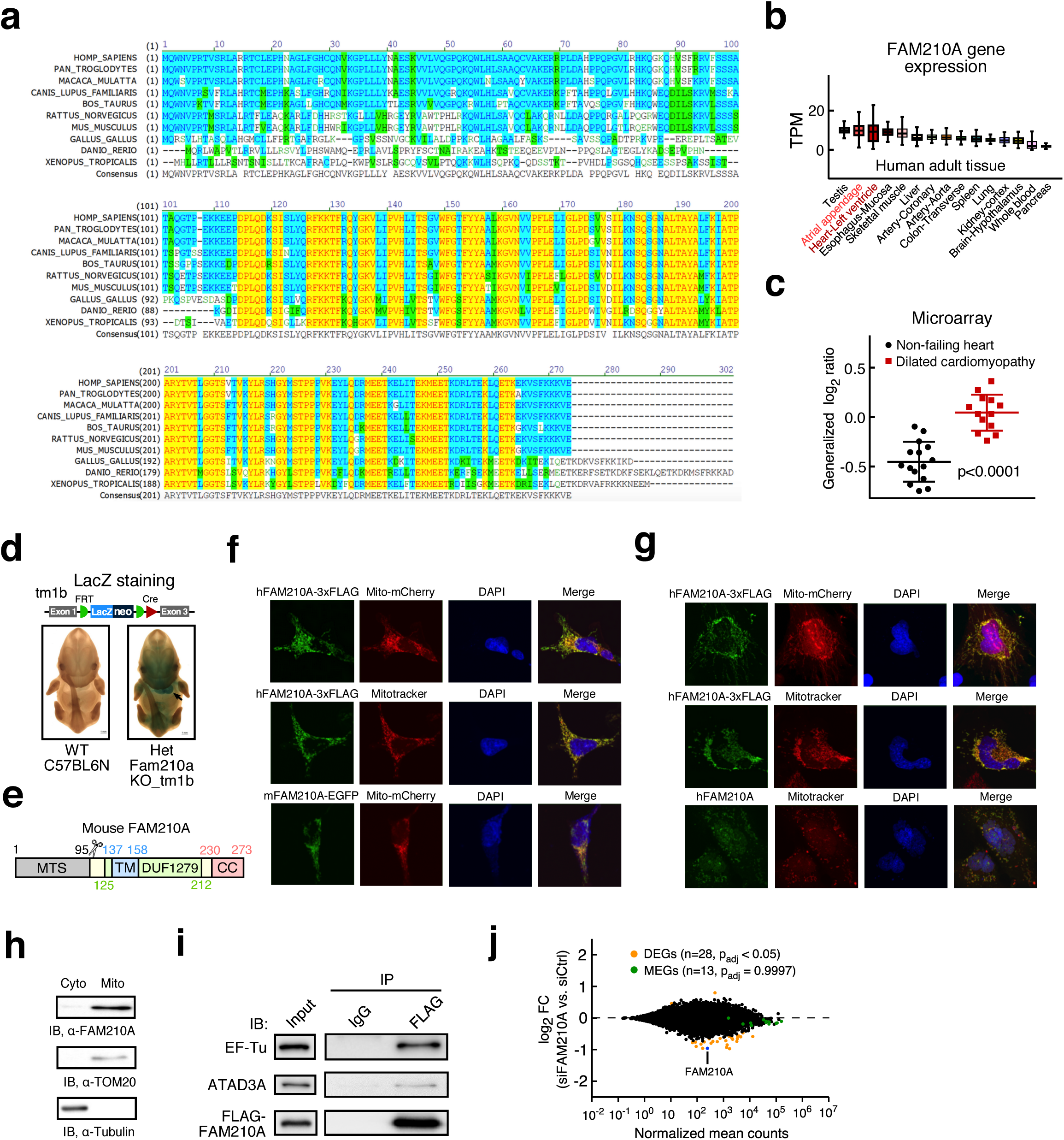
Evolution conservation, organ-specific expression, and cellular localization of FAM210A. **a** Evolutionary conservation of FAM210A protein sequence from 10 representative species. **b** The expression level of *FAM210A* mRNA in 15 human organs from the GTEx portal database. TPM: transcripts per million of reads. **c** *FAM210A* mRNA expression in dilated cardiomyopathy patients (n=13) and non-failing hearts (n=15) from microarray data in GDS2206 GEO profile. The statistical analysis is performed by Student *t* test. **d** Expression of LacZ reporter in E12.5 murine embryo of heterozygous global Fam210a KO mice. Heart-specific high expression of LacZ was observed in all of 9 heterozygous KO mice (IMPC). Tm1b allele: LacZ tagged null allele. **e** The schematic of FAM210A protein domain composition. **f** Cellular localization of recombinant FAM210A in HEK293T cells. FAM210A-3xFlag or FAM210A-GFP expression plasmid was transfected into HEK293T cells for 48 hrs. Anti-FLAG antibody was used for IF. Mito-mCherry or mitotracker were used to label mitochondria. SFB: S-tag, FLAG-tag, and Biotin-tag. **g** Cellular localization of recombinant FAM210A in human AC16 CM cells. FAM210A-3xFlag or FAM210A-GFP expression plasmid was transfected into AC16 cells for 48 hrs. **h** Cellular fractionation and immunoblot for FAM210A protein in AC16 cells. TOM20 and *α*-Tubulin are marker proteins for mitochondria and cytoplasm, respectively. A representative image is shown in replicated experiments. **i** IP-IB validates FAM210A-interacting proteins in HEK293T cell lysates subjected to IP using pre-immune IgG and anti-FLAG antibodies. A representative image is shown in replicated experiments. **j** RNA-Seq analysis of HEK293T cells with siRNA knockdown of FAM210A for 24 hrs. padj, adjusted p-value.

**Supplementary Fig. 7.**
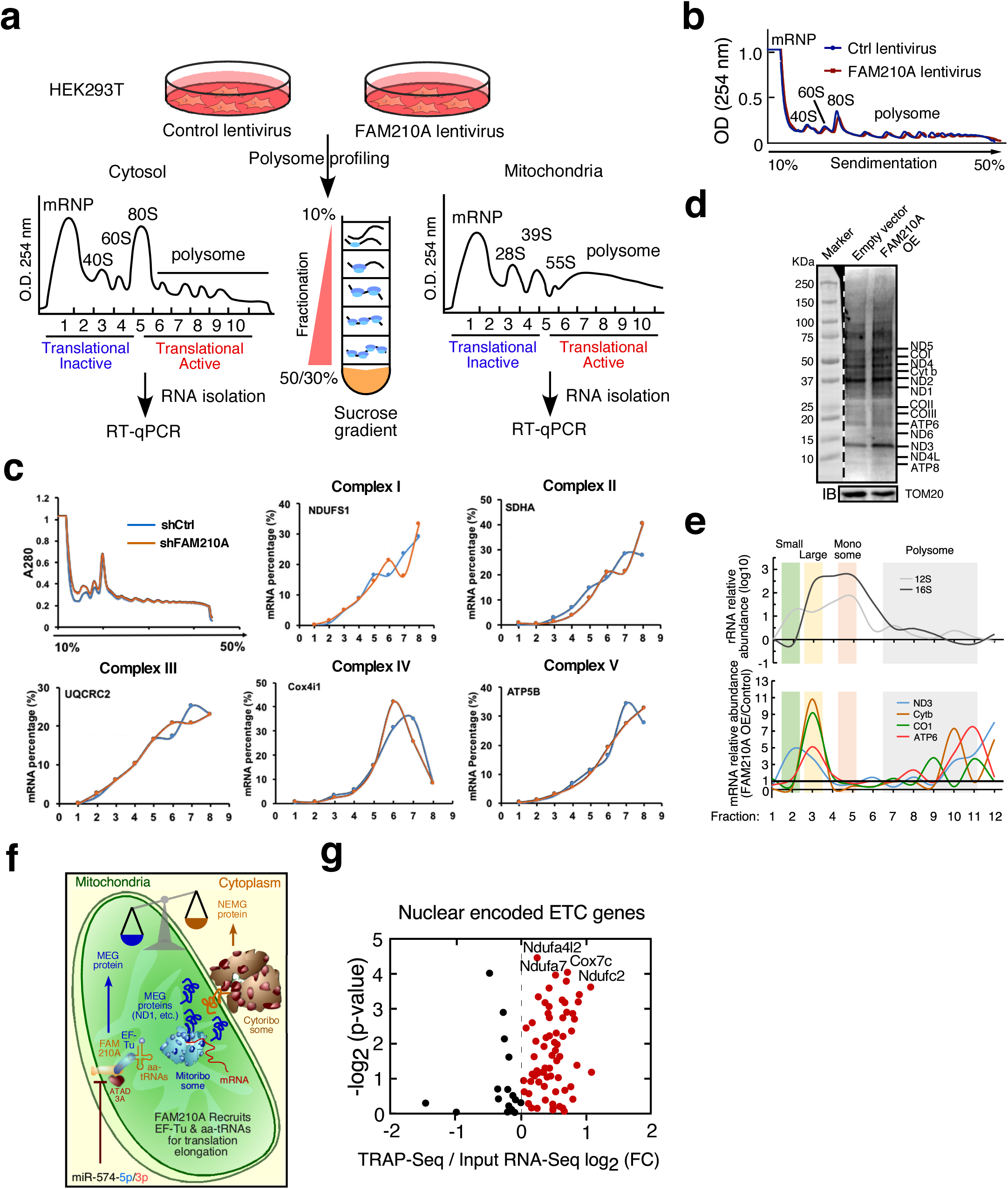
FAM210A influences mitochondrial translation of MEGs but not cytoplasmic translation of NEMGs. **a** The schematic model of polysome profiling assays for measuring the cytoplasmic and mitochondrial translation. **b** Polysome profiling of the cytoplasmic fraction upon overexpression of FAM210A in HEK293T cells. **c** Translation efficiency of multiple NEMGs in HEK293T cells after knockdown of FAM210A. **d** AHA pulse-chase labeling assay for MEGs after lentiviral overexpression of FAM210A. A representative image was shown in replicated experiments. AHA, L-Azidohomoalanine. **e** Mitochondrial polysome profiling of HEK293T cells after stable overexpression of FAM210A followed by RT-qPCR of representative mitochondrial-encoded ETC genes. **f** Working model of ATAD3A-FAM210A-EF-Tu complex in translational control of MEG protein expression in the mitochondria. **g** The translation efficiency of NEMGs in WT murine hearts under TAC surgery. Translation efficiency is calculated by the ratio of TRAP (translating ribosome affinity purification)-Seq divided by RNA-Seq signal.

**Supplementary Fig. 8.**
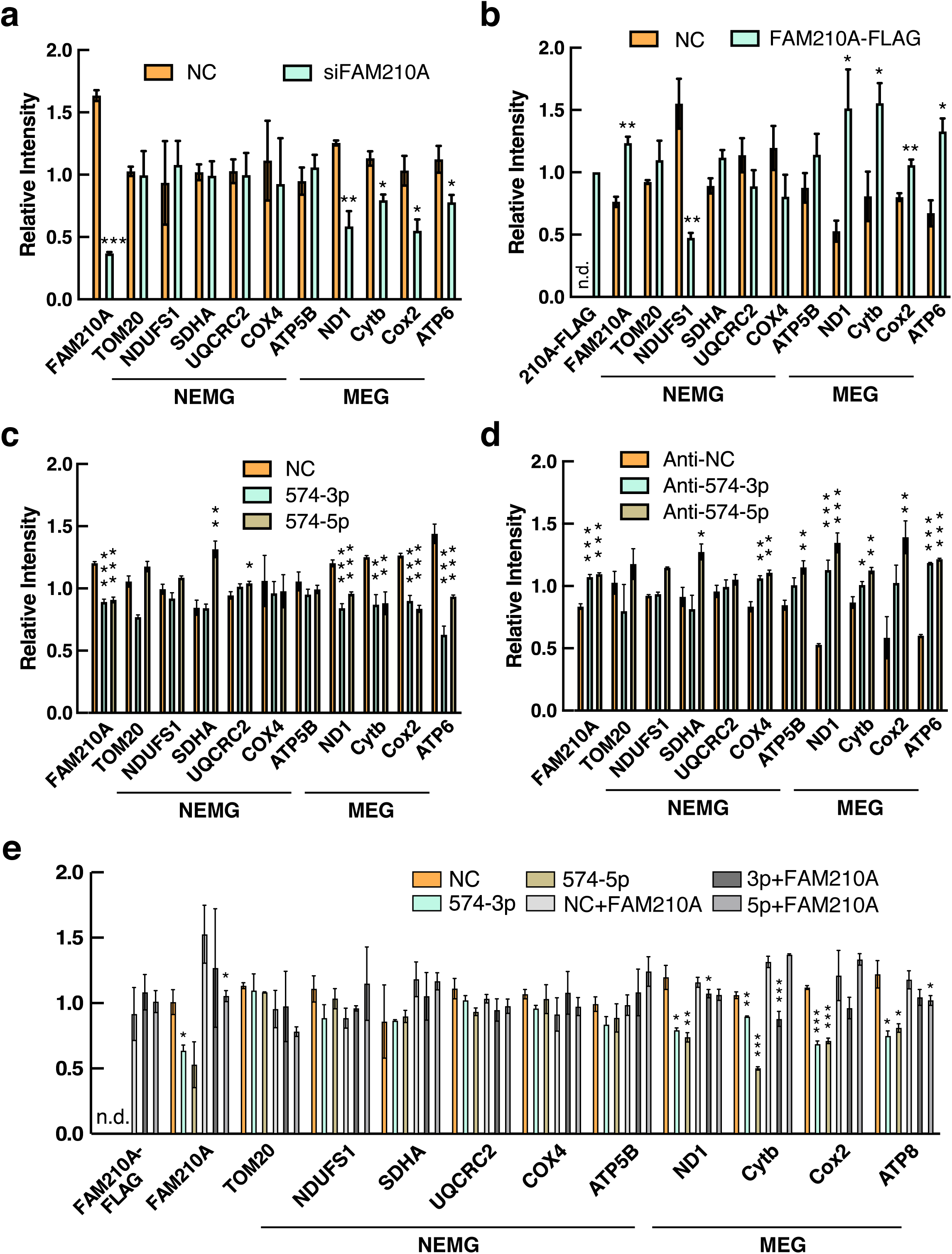
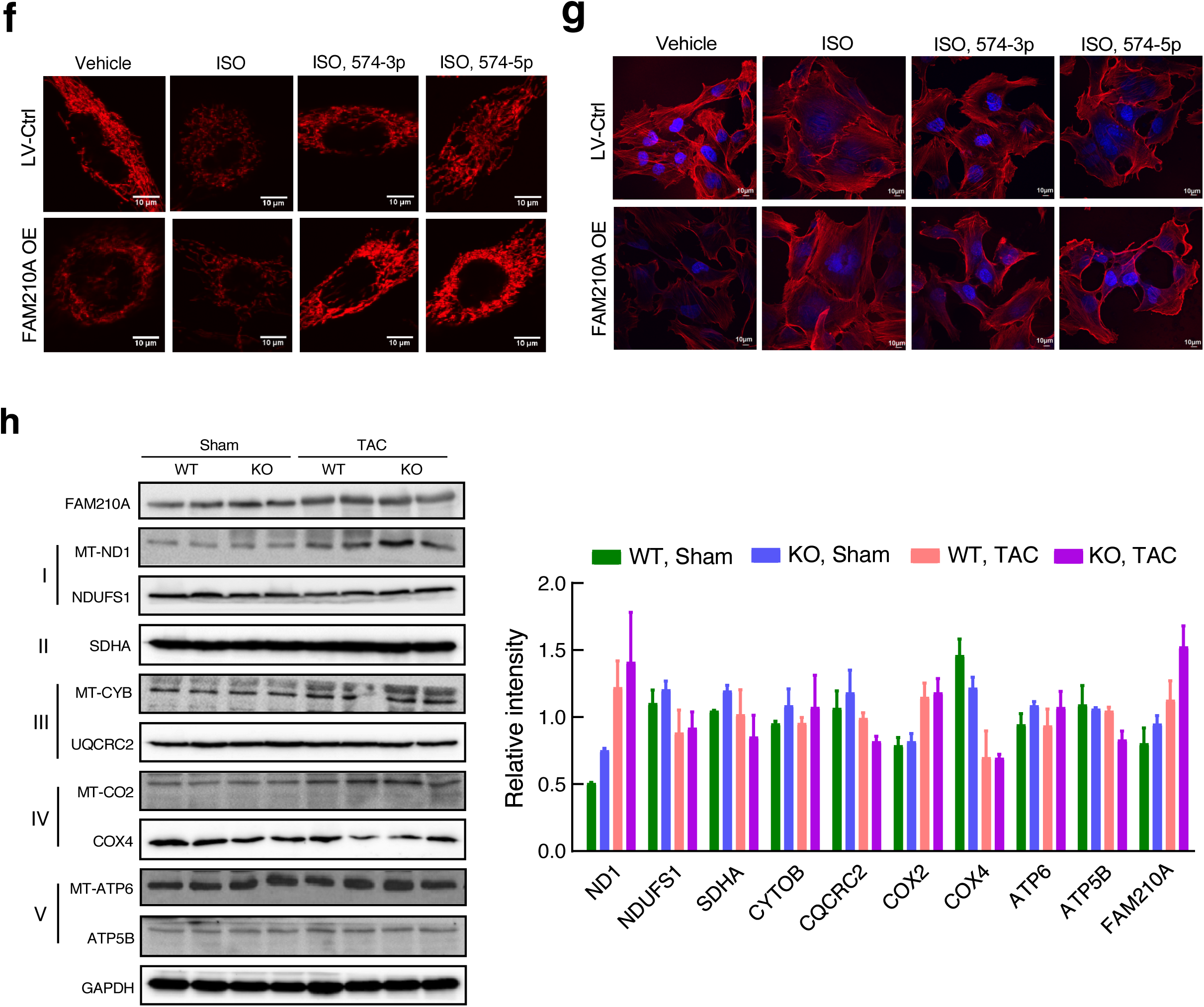
miR-574-FAM210A axis modulates mitochondrial protein expression and mitochondrial activity in AC16 cardiomyocyte cells. **a-e** Quantitative analysis of Western blot data from Fig. 7. n.d., not detected. Each experiment was done in triplicates. **f-g** Representative TMRE and phalloidin staining images of AC16 CM cells. Control AC16 cells and cells with FAM210A stable overexpression were treated with 10 μM ISO for 24 hrs, followed by transfection of 100 nM of miRNA mimics. **h** Protein expression of mitochondrial ETC component genes in WT and miR-574 KO hearts under Sham and TAC surgery (7 days post operation). Images are shown from biologically replicated experiments. Right panel: Quantitative analysis of Western blot data. Data were presented as mean ± SEM. *: p≤0.05, **: p≤0.01, ***: p≤0.001 by one-way ANOVA.

**Supplementary Fig. 9.**
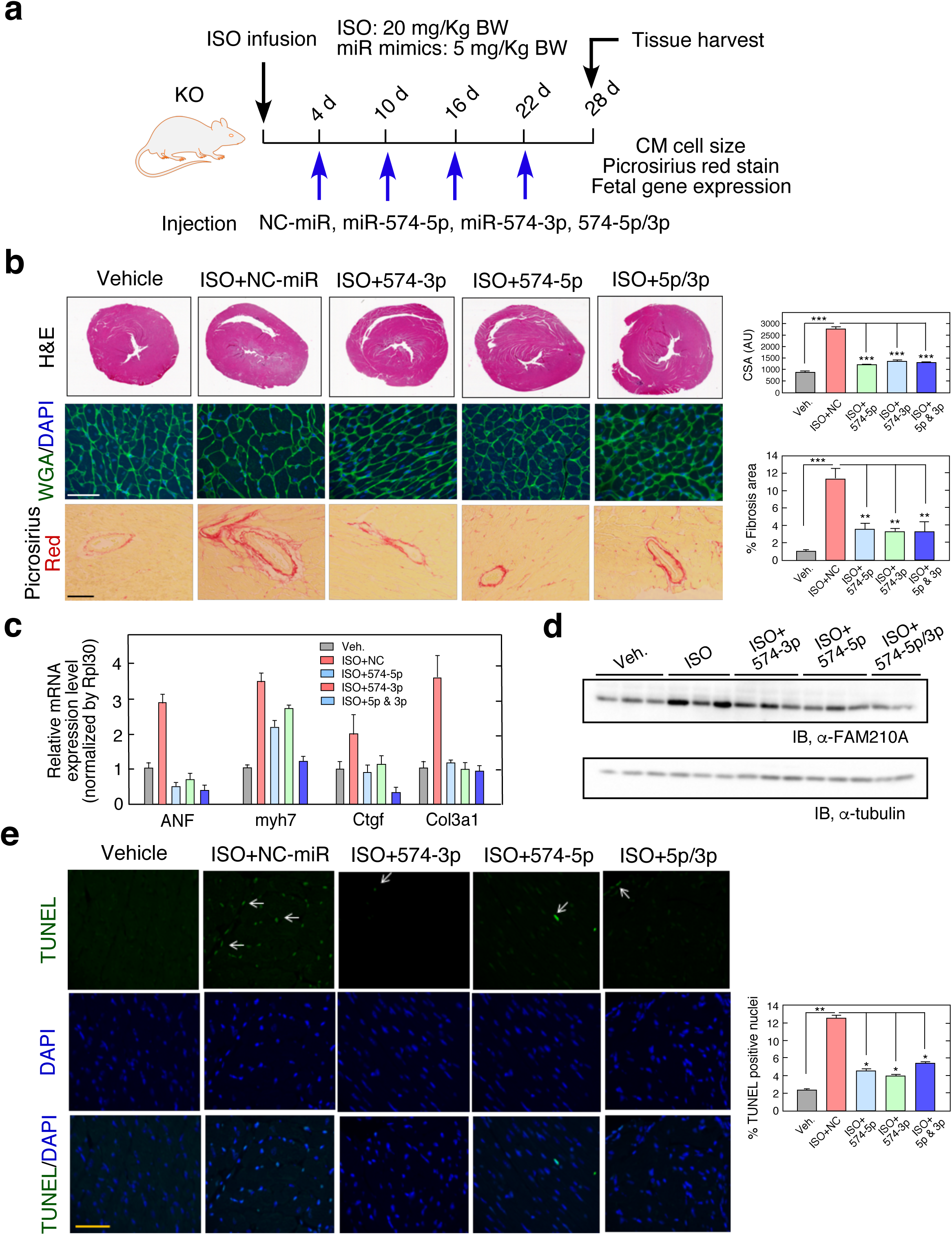
miR-574-5p/3p injection reverses the cardiac pathological remodelling in ISO-treated miR-574 null mice. **a** The schematic of the ISO infusion in miR-574^−/−^ mice using miRNA mimics followed by phenotypic characterizations. HW/TL ratio is measured (n=4-10 per group). NC, negative control. **b** H&E, WGA staining, and picrosirius red staining of murine hearts in the rescue models. **c** RT-qPCR of hypertrophy and fibrosis marker genes in the hearts of miR-574^−/−^ mice with ISO treatment. **d** Western blotting of FAM210A in the hearts of rescue mouse models. **e** TUNEL assay of murine hearts in the rescue models. Data were presented as mean ± SEM. *: p≤0.05, **: p≤0.01, ***: p≤0.001 by one-way ANOVA.

**Supplementary Fig. 10.**
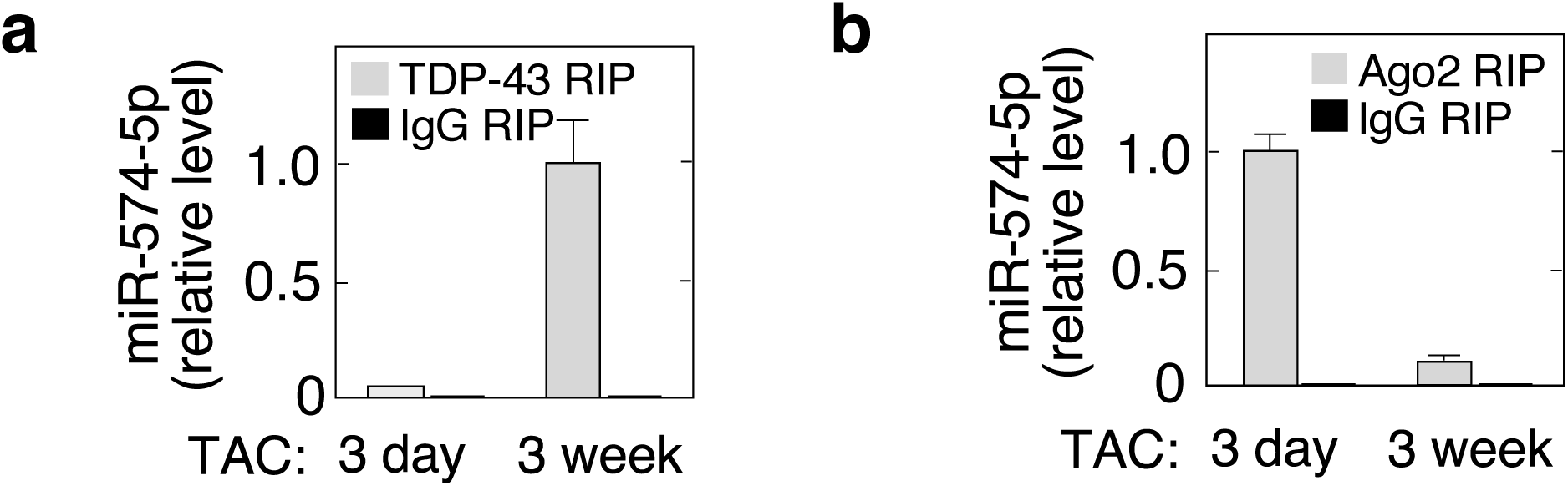
TDP-43 binds to miR-574-5p in the late stage of cardiac remodelling. **a.** RIP-qRT-PCR showed that TDP-43 bound to miR-574-5p in 3 weeks post-TAC. **b** RIP-qRT-PCR showed that interaction of Ago2 with miR-574-5p was reduced in 3 weeks post-TAC.

## ACKNOWLEDGMENTS

We are grateful to Yonggang Zheng, Qiuqing Wang, and Jordan Pappas for critical reading of the manuscript. We appreciate the technical assistance from Ronald Conlon (Case Western Reserve University), Orazio Slivano (Aab CVRI), and Deanne Mickelsen (Aab CVRI) in generating miR-574^−/−^ mice, histology, and surgical operations, respectively. RNA sequencing and primary data analysis were performed by Jason R Myers from the Genomics Research Center at the University of Rochester. Electron microscope imaging is performed by Chad Galloway and Karen de Mesy Bentley (Electron Microscopy Shared Resource Laboratory). None of the authors have any financial conflict of interest related to the research described in this manuscript.

## AUTHOR CONTRIBUTIONS

P.Y. designed the experiments, analyzed the data, wrote the manuscript, and contributed to all subsequent revisions. K.C.V.S and J.W. performed the experiments and analyzed the data. F.J. and O.H. performed some experiments and analyzed the data, A.M. performed echocardiography experiments. K.W. and S.G. performed the mass spectrometry analysis, T.Y. contributed to helpful discussion, manuscript editing, and preparation, W.H.W.T prepared and provided human samples, E.S. and C.Y. contributed to helpful discussion, manuscript editing, and technical support.

## Supplemental discussion

In our IP-mass spectrometry (MS) analysis, EF-Tu is among the top enriched interacting proteins of FAM210A. ATAD3A is located in the mitochondrial inner membrane and has strong interaction with FAM210A detected by anti-ATAD3A antibody IP-MS and quantified by SILAC labelling in the literature (Nucleic Acids Res 2012 40: 6109-21). In the same report, the authors also showed that ATAD3A is associated with EF-Tu (TUFM) in a complex. Here, we used anti-FAM210A antibody to perform IP-MS and identified both EF-Tu and ATAD3A as binding partners of FAM210A in a reversed direction. EF-Tu is known to be mostly associated with the mitochondrial inner membrane (where the mitochondrial ribosomes are located) despite its absence of transmembrane domains (JBC 2007 282: 4076-4084), but the membrane association mechanism has not been discovered. Both ATAD3A and EF-Tu are known to participate in mitochondrial translation. FAM210A may act as a putative adaptor protein to recruit the other two proteins to the membrane since it is a single transmembrane protein localized in the mitochondrial inner membrane. Therefore, we focus on its binding partners in the translation machine. We have confirmed the interaction of FAM210A with EF-Tu and ATAD3A (Fig. 6h, j) and mapped the interacting domain in FAM210A (Fig. 6i). We have also provided evidence to support the regulatory function of FAM210A in MEG protein expression using multiple methods (Fig. S7d,e, Fig. 7a-e, and Fig. S8a-e). The ideal method is to reconstitute the translation machinery in vitro by adding or removing FAM210A from the system to test its detailed molecular mechanism in regulating mitochondrial translation. However, to date, there is no available in vitro translation system to perform this type of assay (FEBS Lett. 2014 588: 2496-2503; Trends Biochem Sci. 2017 42:625-639; Cell Tissue Res. 2017 367:5-20). Therefore, we propose a hypothetical working model in Fig. S7f and we will test this hypothesis and study the detailed mechanism in our future work by blocking the interaction between FAM210A and EF-Tu or ATAD3A via mutating potential FAM210A interacting domains followed by measurement of MEG translation using polysome profiling.

In a recent report, the authors performed RNA-Seq in tamoxifen-inducible skeletal muscle-specific Fam210a conditional knockout mouse-derived cells at the late stage and identified differential expression of a number of muscle cell differentiation-related genes. Therefore, it is possible that FAM210A may be involved directly or indirectly in regulating muscle protein functions. We used crude mitochondrial lysate to perform the IP-MS analysis, and we have not validated the FAM201A-bound cytosolic muscle proteins are authentic interacting partners or abundant contaminant proteins. We observed minor localization of FAM210A in the outer membrane, which may interact with cytosolic muscle proteins for potential crosstalk between mitochondria and cytoskeleton system. We have also noticed that there is another major type of interacting partners for FAM210A, metabolic enzymes, in addition to translation machine component proteins. We cannot rule out the possible role of FAM210A in mitochondrial metabolism as a multi-tasking adaptor molecule. This needs to be thoroughly and systematically analyzed in the future using cardiomyocyte-specific Fam210a conditional knockout mouse. We think that gene expression changes need to be also measured at the early stage of disease progression prior to any severe symptoms occur in order to dissect the direct downstream effector molecules or pathways of FAM210A without getting complicated by secondary effects. We are currently working on a Myh6-Cre driven tamoxifen-inducible Fam210a conditional knockout mouse and generating a cell type (e.g., myocyte) specific Fam210a-overexpressing transgenic mouse. We expect that we will be able to fully address the biological function and molecular mechanism of FAM210A using these two mouse models in the future.

## SUPPLEMENTAL MATERIAL

### Supplemental Methods

#### Plasmids, antibodies, miRNAs, siRNAs, and qPCR primers

Full-length coding sequence (CDS) of human FAM210A was cloned into pcDNA3.1(+) with 3x Flag in the C-terminus of the genes. Full-length CDS of mouse Fam210a was cloned into pEGFP-N1 with EGFP in the C-terminus of the gene. The primer sequences for cloning FAM210A gene are as follows:

FAM210A-3xFlag-F (BamHI): 5’ CGGGATCCCAAAATGCAATGGAATGTACCACG 3’

FAM210A-3xFlag-R (EcoRI): 5’ CGGAATTCTTCCACTTTTTTCTTAAAGGAAAC 3’

Fam210a-EGFP-F (XhoI): 5’ CCGCTCGAGCAAAATGCAATGGAATGTACCAC 3’

Fam210a-EGFP-R (EcoRI): 5’ CGGAATTCGTTCCACTTTTTTCTTAAAGGAC 3’

The 2^nd^ generation of lentivirus package vectors, psPAX2 and pMD2.G, were gifts from Dr. Guang Yang at City of Hope National Medical Center. pLV-CMV-EF1-GP vector was used for the construction of FAM210A overexpression plasmid. Lentivirus-based shRNA vectors against mouse (vector name: pLV[shRNA]-EGFP:T2A:Puro-U6>mFam210a[shRNA#1]) and human (vector name: pLV[shRNA]-EGFP:T2A:Puro-U6>hFAM210A[shRNA#1]) FAM210A were designed and cloned through VectorBuilder website, and purchased from the company.

Primary antibodies used in this study include: rabbit polyclonal anti-FAM210A antibody (HPA014324, validated by ProteinAtlas) and mouse monoclonal anti-Flag antibody (F3165) were purchased from Sigma-Aldrich. Mouse monoclonal antibodies against VDAC (sc-390996), ATP5B (sc-74549), TOM20 (sc-17764), EF-Tu (sc-393924), ATAD3 (sc-376185), NDUFS1(sc-271510), COX4 (sc-376731), COX2 (sc-514489), NDUFA8 (sc-398097), SDHA (sc-166909), and UQCRC2 (sc-390161) were obtained from Santa Cruz Biotechnology, Inc. Rabbit polyclonal antibodies against Flag tag (20543-1-AP), *α*-Tubulin (11224-1-AP) and ND1 (19703-1-AP), and mouse monoclonal anti-GAPDH (60004-1-lg) were from Proteintech. Rabbit polyclonal antibodies against CYTB (PA5-43533), ATP8 (PA5-68103) and ATP6 (PA5-37129) were from ThermoFisher Scientific. Rabbit monoclonal anti-*β*-actin (13E5, #4970) was from Cell Signaling Technology.

The *mir*Vana® miRNA mimics are chemically modified double-stranded RNAs designed for *in* vivo injection studies in mouse models as well as *in vitro* transfection in cell culture studies (designed and synthesized by ThermoFisher Scientific). Scrambled *mir*Vana® miRNA mimics (Cat. No. 4464061) were used as negative control. The mirVana miRNA mimics for mmu-miR-574-5p (Cat. No. 4464070), miR-574-3p (Cat. No. 4464066), and Silencer Select siRNA against mouse (Cat. No. 173500) or human (Cat. No. 129436) were used. For inactivation of endogenous miRNAs, we use Anti-miR™ miRNA Inhibitor Negative Control #1 (Cat. No. AM17010), Anti-miR™ miR-574-5p Inhibitor (Cat. No. AM17010, assay ID AM13081), and Anti-miR™ miR-574-3p Inhibitor (Cat. No. AM17010, assay ID AM12848). FAM210A were purchased from ThermoFisher Scientific. miRNAs were transfected into cells using lipofectamine 3000 (ThermoFisher Scientific) following the manufacturer’s instructions. For AC16 cell transfection, 100-150 nM miRNA mimics (scrambled miRNA, miR-574-5p, or miR-574-3p) or anti-miR inhibitors (NC anti-miR, anti-miR-574-5p, or anti-miR-574-3p) were transfected for 24 hrs before measuring hypertrophic growth of the cells. For HEK293T cell transfection, 100 nM miRNA mimics (scrambled miRNA, miR-574-5p, or miR-574-3p) were transfected for 24 hrs before measuring expression of ETC component proteins. 100 nM scrambled siRNA or FAM210A siRNA and 1 μg of FAM210A overexpression plasmid were transfected for 24 hrs before measuring expression of ETC component proteins.

Total RNA was isolated from the mouse and human left ventricle samples using Trizol reagent (ThermoFisher Scientific). For RT-qPCR detection of miRNA expression, 1 μg of RNA was used for reverse transcription using the miScript II RT Kit (Qiagen, Cat. No. 218161). 1 μl of cDNA and miScript® SYBR® Green PCR kit (Cat. No. 218073) were used for quantitative PCR. qPCR primers were purchased from miScript Primer Assays (Qiagen): Hs_miR-574-5p_2 miScript Primer Assay (Cat. No. MS00043617), Hs_miR-574-3p_1 miScript Primer Assay (Cat. No.: MS00032025), Hs_SNORD68_11 miScript Primer Assay (Cat. No. MS00033712), Hs_RNU6_11 miScript Primer Assay (Cat. No. MS00033740). For mRNA reverse transcription, iScript^TM^ cDNA Synthesis Kit (Bio-Rad, Cat. No. 1708891) was used. iTaq™ Universal Probes Supermix (Bio-Rad, Cat. No. 1725134) and TaqMan Real-time PCR assays (ThermoFisher Scientific) was used for TaqMan qPCR assay. Taqman qPCR assay probes (mouse) were purchased from ThermoFisher Scientific and the Cat. numbers are listed as follows: Fam114a1 (4351372, Mm00471781_m1), ANF (Nppa, Mm01255747_g1), BNP (Nppb, Mm01255770_g1), Myh6 (Mm00440359_m1), Myh7 (Mm01319006_g1), Col1a1 (Mm00801666_g1), Col1a2 (Mm00483911_g1), Col3a1 (Mm00802305_g1), Ctgf (Mm01192932_g1), Rpl30 (Mm01611464_g1), Gapdh (Mm99999915_g1). iTaq Universal SYBR Green Supermix (Bio-Rad, Cat. No. 1725125) and qPCR primers (synthesized from IDT) were used for SYBR green qPCR assay. The sequences of SYBR qPCR primers are listed as below (h: human, m: mouse, h/m: human and mouse).

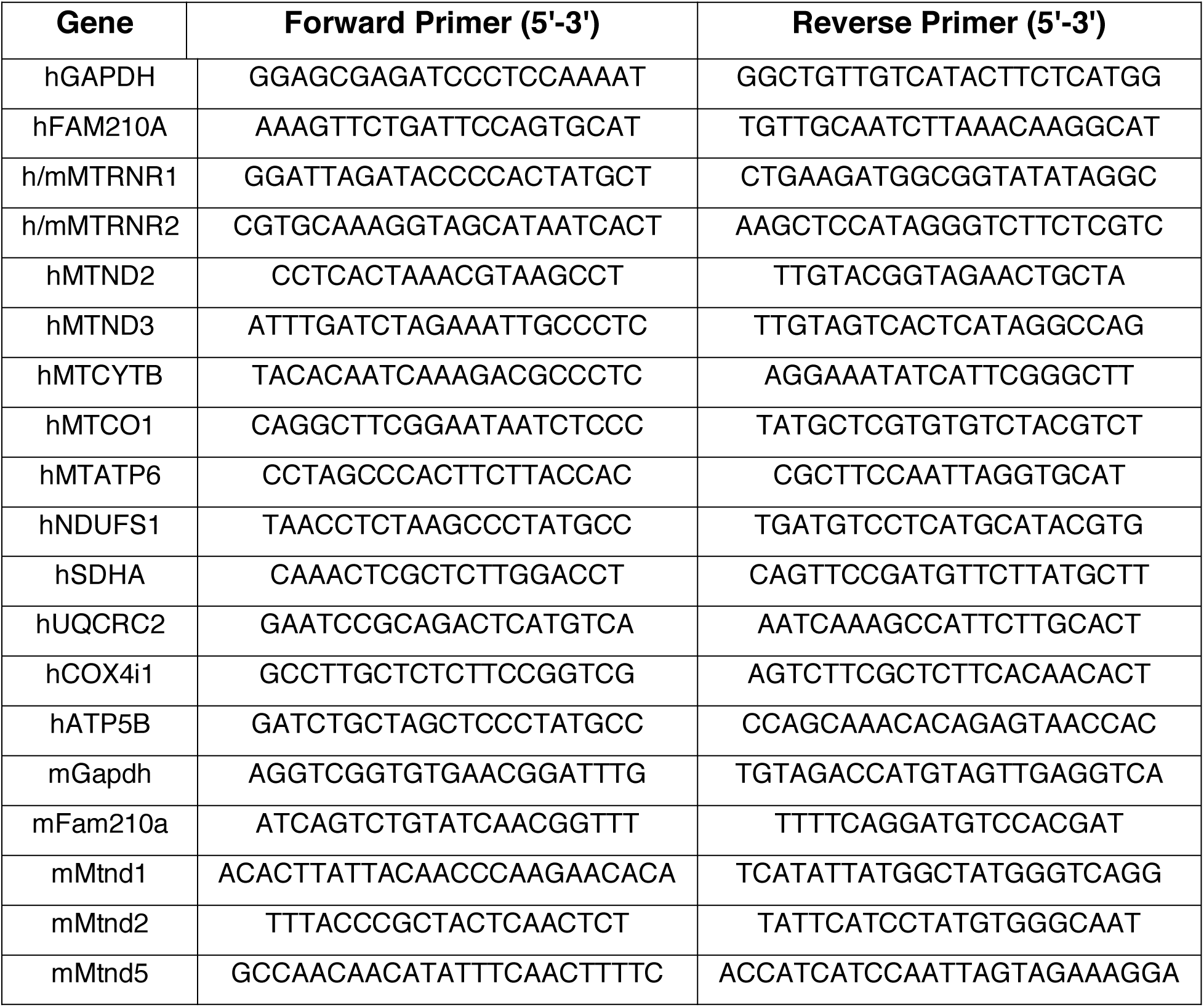
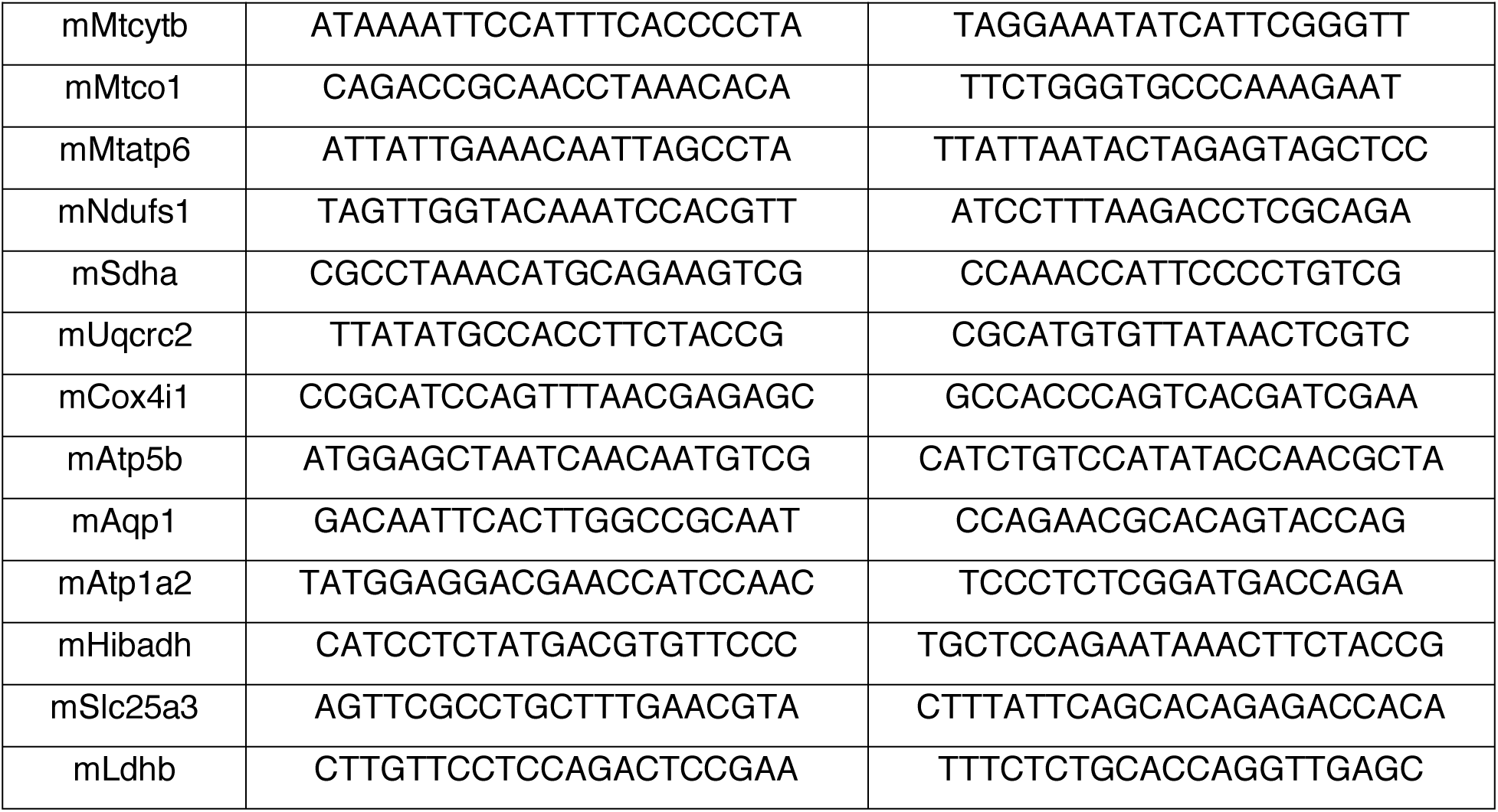

#### Mice

Three Sanger MirKO ES cell lines for Mir574 (produced by Wellcome Trust Sanger Institute)^1^ were purchased from the mutant mouse regional resource center (MMRRC). miR-574^−/−^ global knockout chimera founder mice were generated in the Case Western Reserve University Transgenic Core Facility (Dr. Ronald Conlon). We generated a miR-574 targeted male chimera mouse in the C57BL/6J background, performed germline transmission, and backcrossed the mice to C57BL6J mice (purchased from Jackson Laboratory) for more than 10 generations. A 182-bp DNA region was deleted including the microRNA574 gene (96-bp for pre-miR-574 sequence) in the intron 1 of the host gene Fam114a1. Puromycin resistance cassette was removed by breeding with C57BL/6-Tg(Zp3-cre)93Knw/J mouse (ZP3-Cre transgene), which expresses Cre recombinase in oocytes, resulting in a null allele. For experiments with miR-574^−/−^ mice, control mice of the same age and gender from littermates or sibling mating were used. All animal procedures were performed in accordance with the National Institutes of Health (NIH) and the University of Rochester Institutional guidelines. Genotyping PCR and electrophoresis protocols are shown as below (miR-574 KO mice and genotyping protocols were donated to MMRRC):

| Reagent/Constituent | Volume (μL) |
| --- | --- |
| Water | 17.5 |
| 10x Buffer | 2.5 |
| MgCl<sub>2</sub> (stock concentration is 25 mM) | 1 |
| dNTPs (stock concentration is 10 mM) | 1 |
| Primer 1 (stock concentration is 20 μM) | 0.25 |
| Primer 2 (stock concentration is 20 μM) | 0.25 |
| Taq Polymerase 5 Units/μL | 0.5 |
| DNA (50-200 ng/μL) | 2 |
| **TOTAL VOLUME OF REACTION:** | **25 μL** |

| Steps | Temp ($^{\circ}$C) | Time (m:ss) | # of Cycles |
| --- | --- | --- | --- |
| 1. Initiation/Melting | 94 | 5:00 | 1 |
| 2. Denaturation | 94 | 0:30 |  |
| 3. Annealing | 56 | 0:30 | 34x |
| 4. Elongation | 68 | 0:30 |  |
| 5. Amplification | 68 | 5:00 | 1 |
| 6. Finish | 4 | ∞ | n/a |

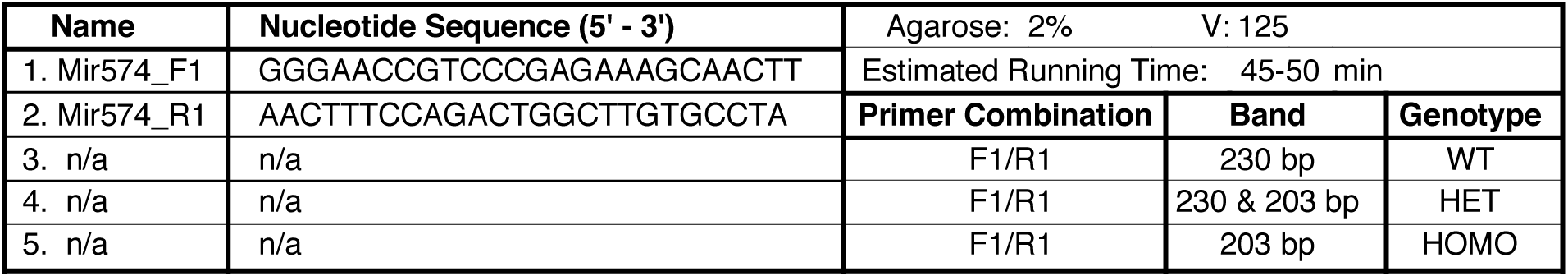

#### *In vivo* therapeutic model using miRNA mimics

Based on the recommendation from the nanoparticle user instruction, 75-100 μg miRNA with chemical modification (resistant to nuclease degradation in vivo) needs to be used per injection. In our experiments, miRNA mimics were used at the dose of 5 mg/Kg (high dose) or 1 mg/Kg (low dose) body weight in the volume of 150-200 μl for injections in male WT C57BL/6J mice (10-12 weeks old, under TAC surgery) or miR-574 null mice (10-12 weeks old, with ISO infusion). The *mir*Vana® miRNA mimics for miR-574-5p/3p and scrambled *mir*Vana® miRNA mimics (negative control) (∼100 μg) were dissolved in ∼150-200 μl RNase-free water. The diluted miRNA mimics were incubated with 50 μl of nanoparticle-based in vivo transfection reagent (Altogen Biosystems, Cat. No. 5031) in sterile tubes for 20 mins at RT. Transfection enhancer (10 μl, Altogen Biosystems, Cat. No. 1799) was added to the mixture, vortexed gently, and incubated for 5 mins at RT. The nanoparticle-miRNA mimics complex was mixed with an appropriate volume of the sterile solution of 5% glucose (w/v), and delivered into the murine heart by intravenous tail vein injections (after mice are anesthetized using 2.0% isoflurane) once a week after TAC surgery (or 4 days after ISO infusion starts) following the manufacturer’s instruction and a previous report.^2^

#### Echocardiography

The echocardiographic image collection is performed using a Vevo2100 echocardiography machine (VisualSonics, Toronto, Canada) and a linear-array 40 MHz transducer (MS-550D). Image capture is performed in mice under general isoflurane anesthesia with heart rate maintained at 500-550 beats/min. LV systolic and diastolic measurements are captured in M-mode from the parasternal short axis. Fraction shortening (FS) is assessed as follows: % FS = (end diastolic diameter - end systolic diameter) / (end diastolic diameter) x 100%. Left ventricular ejection fraction (EF) is measured and averaged in both the parasternal short axis (M-Mode) using the tracing of the end diastolic dimension (EDD) and end systolic dimension (ESD) in the parasternal long axis: % EF=(EDD-ESD)/EDD. Hearts are harvested at multiple endpoints depending on the study.

#### Langendorff procedure for adult mouse cardiomyocyte/fibroblast isolation

<u>Preparation:</u> 10x perfusion buffer (pH 7.4) is prepared and stored at 4°C the day before the procedure. Coverslips are sterilized with 70% EtOH for 15 mins, then rinsed with 100% EtOH, and air-dried overnight. On the day of operation, TC plates are coated with laminin for cardiomyocytes (CMs) for >2 hrs at 37°C. The perfusion apparatus is turned on, warmed up, and washed by Milli Q H2O, 70% EtOH, Milli Q H2O, and rinsed with 200 ml Milli Q H2O before first Langendorff of the day. Fresh perfusion buffer (PB) and digestion buffer (DB) are made and sterilized through 0.2 μm filters. The perfusion apparatus is filled with 1x PB, and 10 cm petri dish is filled with 1x PB to place heart before cannulation. Then the empty 15-ml Falcon with 100 μm mesh is prepared.

<u>Protocol:</u> Mice are anesthetized with anesthetic solution (0.5 ml ketamine [100 mg/Kg]/ xylazine [10 mg/Kg]/PBS + 0.2 ml 1000 U/ml heparin), and sacrificed by cervical dislocation. Hearts are removed, cannulated and perfused with 1x PB for 4 mins, switched to DB for 3 mins, and then perfused with 32 ml DB with CaCl2 for 8 mins. Hearts are then released from cannula and atria, and great vessels are removed. Ventricles are placed in sterile 35 mm dish with 2.5 ml DB and shredded into several pieces with forceps. 5 ml stopping buffer (SB) is added and pipetted several times until tissues disperse readily, and solution turns cloudy. The cell solution is transferred to a conical tube with 100 μm mesh to filter cells from tissue. Dishes are rinsed with 2.5 ml SB and added to a tube through a mesh (final volume is 10 ml). CMs are settled by incubating the tube at 37°C for 15 mins. The supernatant (containing cardiac fibroblasts) is transferred to a fresh tube (∼8 ml), and 10 ml SB are added to the pellet of CMs. CMs are treated with Ca^2+^ ramp, swirled to mix every 2-3 mins, followed by adding 10 μl 100 mM Ca^2+^ for 2 mins, 40 μl 100 mM Ca^2+^ for 4 mins, and 90 μl 100 mM Ca^2+^ for 7 mins. The laminin is removed from plates and CMs are added in final SB with Ca^2+^. Plates are centrifuged for 5 mins at 1,000 rpm at 4°C, allowing CMs to sit at 37°C for ∼1 hr in SB, and then washed 3 times with 2 ml 1x PBS. CMs are washed for 3 wells at a time by swirling 3 times, tilted, aspirated, and replaced with 1x PBS. One adult heart should yield at least 2-3 full 6 well plates of CMs. We aim for less than 10% dead and round CMs. Cardiac fibroblasts (CFs) from the supernatant are pelleted for 5 mins at 1000 rpm at 4°C. CFs are plated in 4-5 ml CF-specific media in 60 mm plate. CFs are washed vigorously 3-5 times with 2 ml 1x PBS after 2-3 hrs and replaced with fresh CF media.

Solutions for Langendorff operations:

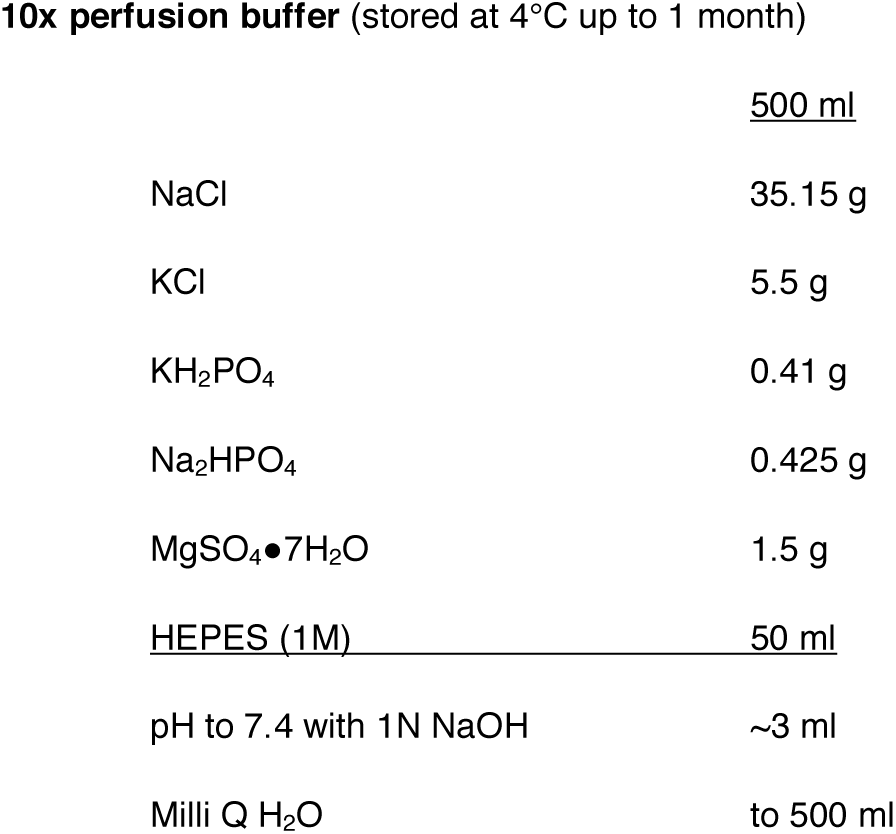
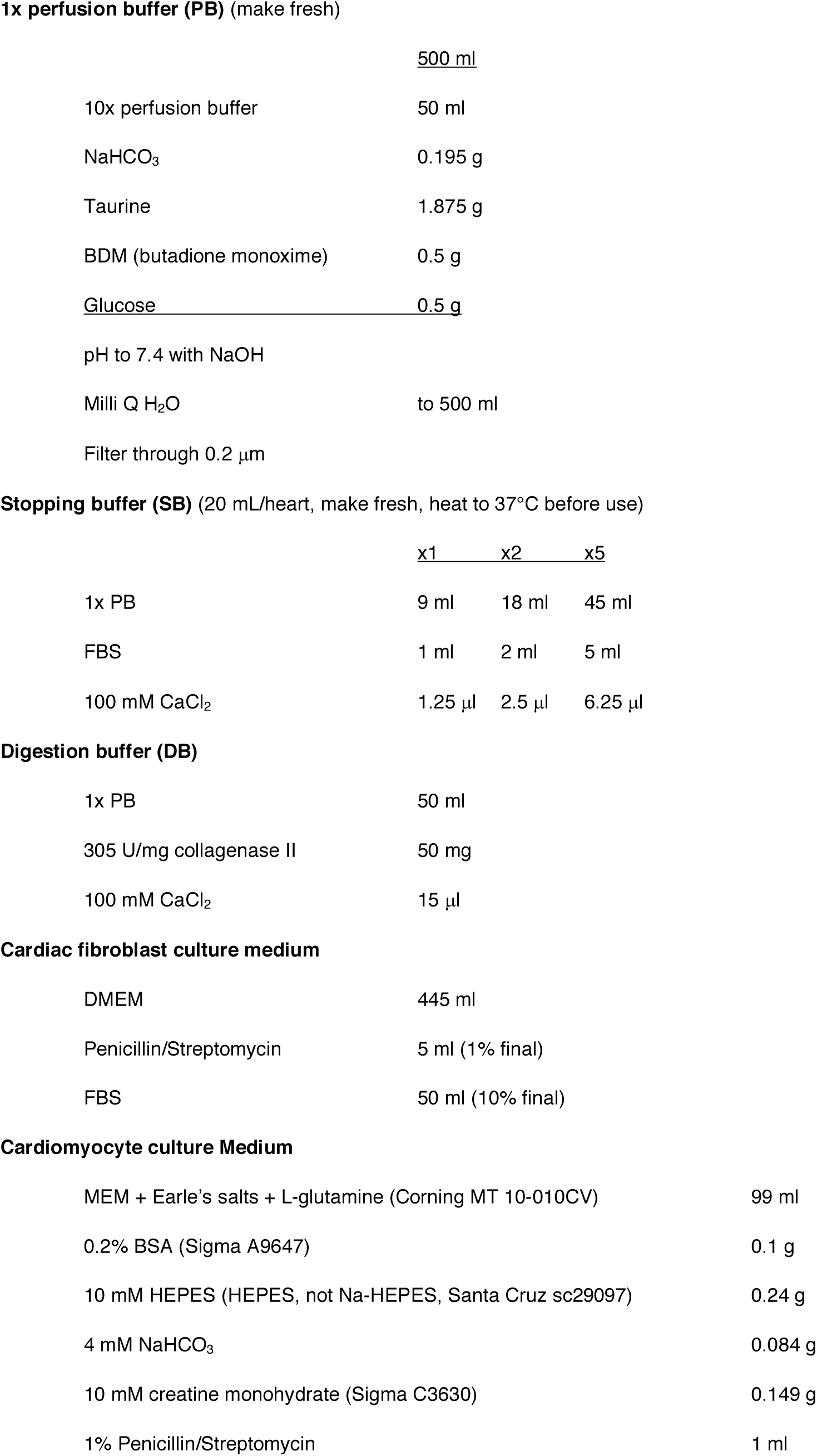
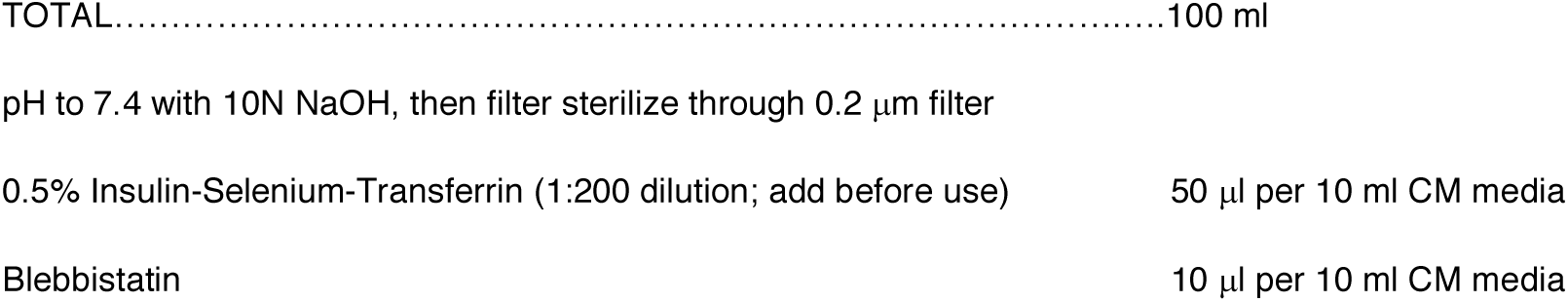

|  | <u>500 ml</u> |
| --- | --- |
| NaCl | 35.15 g |
| KCl | 5.5 g |
| $\text{KH}_2\text{PO}_4$ | 0.41 g |
| $\text{Na}_2\text{HPO}_4$ | 0.425 g |
| $\text{MgSO}_4 \cdot 7\text{H}_2\text{O}$ | 1.5 g |
| <u>HEPES (1M)</u> | <u>50 ml</u> |
| pH to 7.4 with 1N NaOH | ~3 ml |
| Milli Q $\text{H}_2\text{O}$ | to 500 ml |
| **1x perfusion buffer (PB)** (make fresh) |  |
|  | <u>500 ml</u> |
| 10x perfusion buffer | 50 ml |
| NaHCO<sub>3</sub> | 0.195 g |
| Taurine | 1.875 g |
| BDM (butadione monoxime) | 0.5 g |
| <u>Glucose</u> | <u>0.5 g</u> |
| pH to 7.4 with NaOH |  |
| Milli Q H<sub>2</sub>O | to 500 ml |
| Filter through 0.2 µm |  |

**Stopping buffer (SB)** (20 mL/heart, make fresh, heat to 37°C before use)
|  |  |  |  |
| --- | --- | --- | --- |
|  | <u>x1</u> | <u>x2</u> | <u>x5</u> |
| 1x PB | 9 ml | 18 ml | 45 ml |
| FBS | 1 ml | 2 ml | 5 ml |
| 100 mM CaCl<sub>2</sub> | 1.25 µl | 2.5 µl | 6.25 µl |
| **Digestion buffer (DB)** |  |
| 1x PB | 50 ml |
| 305 U/mg collagenase II | 50 mg |
| 100 mM CaCl<sub>2</sub> | 15 µl |
| **Cardiac fibroblast culture medium** |  |
| DMEM | 445 ml |
| Penicillin/Streptomycin | 5 ml (1% final) |
| FBS | 50 ml (10% final) |
| **Cardiomyocyte culture Medium** |  |
| MEM + Earle's salts + L-glutamine (Corning MT 10-010CV) | 99 ml |
| 0.2% BSA (Sigma A9647) | 0.1 g |
| 10 mM HEPES (HEPES, not Na-HEPES, Santa Cruz sc29097) | 0.24 g |
| 4 mM NaHCO<sub>3</sub> | 0.084 g |
| 10 mM creatine monohydrate (Sigma C3630) | 0.149 g |
| 1% Penicillin/Streptomycin | 1 ml |
| TOTAL..... | 100 ml |
| pH to 7.4 with 10N NaOH, then filter sterilize through 0.2 µm filter |  |
| 0.5% Insulin-Selenium-Transferrin (1:200 dilution; add before use) | 50 µl per 10 ml CM media |
| Blebbistatin | 10 µl per 10 ml CM media |

### Luciferase activity assay

Reporter vectors were generated by inserting the Fam210a 3’UTR fragments containing wild-type or mutated seed sequence into the miRNA reporter vector pmirGlo (Promega). All the primers used to generate the reporter vector were listed as below. The wild-type 3’UTR fragment was amplified and ligated to pmirGlo. Then, 100 ng of wild-type or mutant reporter vectors were co-transfected with 20 nM miRNA (miR-574-5p or miR-574-3p) into HEK293T cells cultured in 24-well plate using lipofectamine 3000 (ThermoFisher Scientific) following the manufacturer’s instruction. Cells were collected at 36 hrs after transfection, and firefly and renilla luciferase activity were detected by Dual-Luciferase Reporter Assay System (Promega).

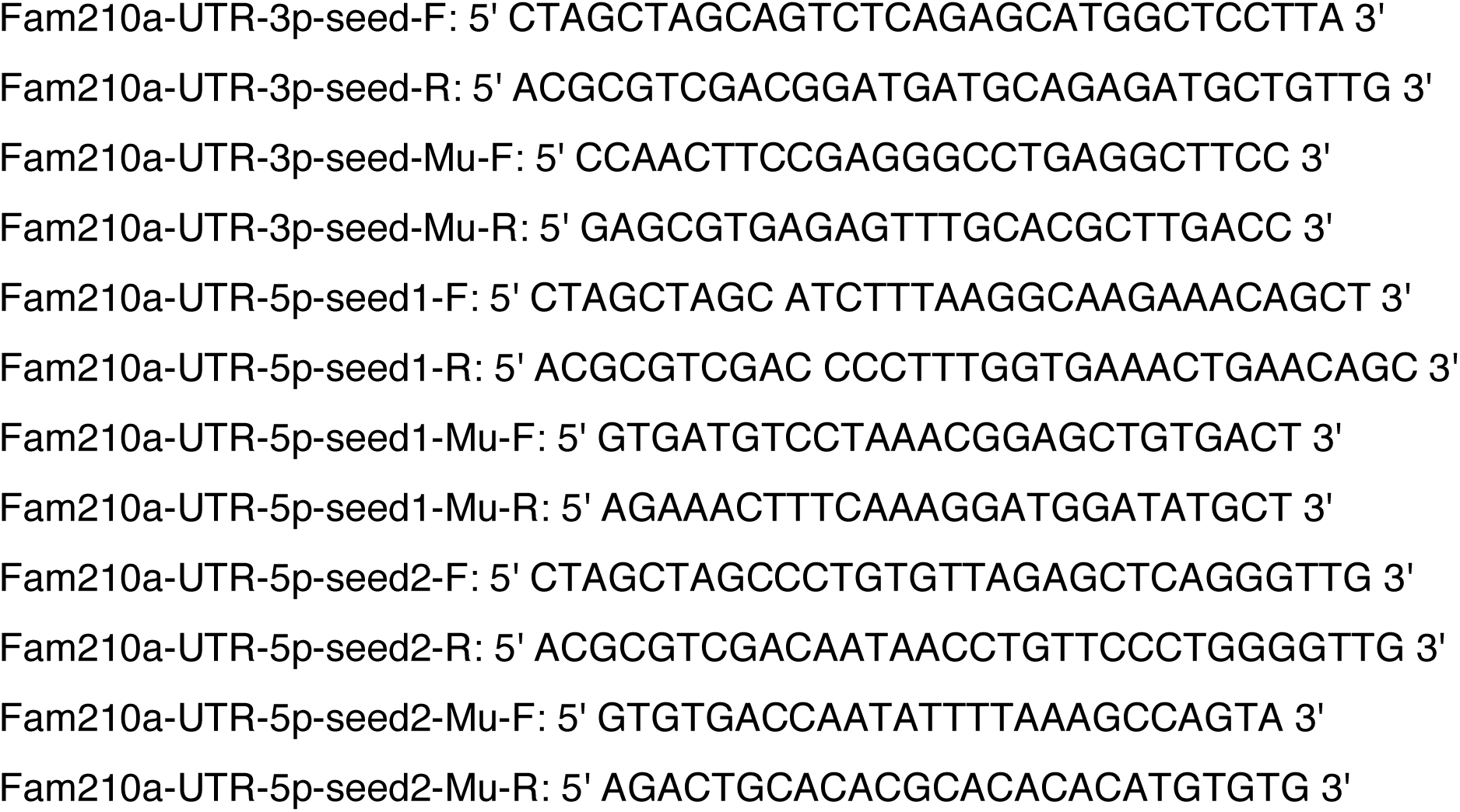

### Mitochondria isolation and mitoplast purification

Mitochondria isolation and mitoplast purification from HEK293T cells and murine hearts were performed as previously decribed^3^. The cells were collected by centrifugation and re-suspended in ice-cold mitochondria isolation buffer (3 mM HEPES-KOH pH7.4, 210 mM mannitol, 70 mM sucrose, 0.2 mM EGTA with protease inhibitor cocktail). Then, the cells were homogenized by a pre-chilled Micro-Tube Homogenizer (Bertin Technologies) for 40-50 strokes. The homogenized cell samples were centrifuged at 500 g for 10 mins to remove the nuclei and unbroken cells. The mitochondria were sedimented by centrifugation at 10,000 g from clear supernatant. To isolate the mitoplast and outer membrane (OM), we used mitochondria isolation buffer containing 0.5 mg/ml digitonin (Sigma) to re-suspend the mitochondria pellet and incubate at 4°C in microtube mixer for 15 mins. The mitoplast was sedimented by 20,000 g centrifugation for 10 mins after adding the equal volume of mitochondria isolation buffer. The pellet contained the matrix and inner membrane (IM) while the supernatant was the OM fraction of the mitochondria. Western blot analysis of OM (TOM20) or IM ETC (ATP5B) protein marker indicates the quality of purification.

### Analysis of cytosolic and mitochondrial translational activity by polysome profiling

Polysome profiling for the nuclear-encoded mitochondrial genes (cytosolic translation) was performed as previously described^4^. Cycloheximide (CHX, 100 μg/ml, Sigma) was added to the cells for 15 mins before lysis to freeze ribosomes on mRNAs in the elongation phase. Around 10^7^ cells were lysed in TMK lysis buffer (10 mM Tris-HCl pH 7.4, 100 mM KCl, 5 mM MgCl2, 1% Triton X-100, 0.5% Deoxycholate, 2 mM DTT) containing 100 μg/ml CHX, 4 U/ml RNase inhibitor (NEB), and proteinase inhibitor cocktail (Roche) on ice for 15 mins. Equal amounts of A260 absorbance from each sample were loaded onto a 10-50% sucrose gradient solution, and centrifuged at 29,000 rpm for 4 hrs; 22 translation fractions were collected from each sample by Density Gradient Fractionation System (BRANDEL). Based on the UV absorbance curve, the 22 fractions were pooled into 8 samples including free mRNP, 40S small ribosome subunit, 60S large ribosome subunit, 80S monosome, light polysomes (di-ribosome, tri-ribosome, etc.), and heavy polysomes (>5 ribosomes). Total RNA was extracted from the same volume of each pooled fraction with Trizol LS (ThermoFisher Scientific), and Renilla luciferase mRNA from *in vitro* transcription was used as RNA spike-in and loading control for RT-qPCR. The mitochondrial polysome profiling was performed as previously described.^5^ In contrast with the cytosolic polysome profiling, the mitochondria were isolated from cells and lysed in TMK lysis buffer. Then the samples were loaded onto a 10-30% sucrose gradient solution and centrifuged following the same procedure as with cytosolic polysome profiling. Twelve pooled fractions were collected from mitochondrial polysome profiling for further analysis of the translation of mitochondrial-encoded genes. The association of mitochondrial-encoded ETC gene mRNAs was analyzed by RT-qPCR. 12S and 16S rRNAs were used for RT-qPCR as quality controls for polysome profiling.

### Interactome capture by Immunoprecipitation-Mass spectrometry (IP-MS)

1. <u>Immunoprecipitation:</u> The total mitochondria were isolated from the whole heart of wild-type mice following the manual using the Mitochondria Isolation Kit for Tissue (Pierce). The mitochondria pellets were re-suspended in NP-40 lysis buffer (50 mM HEPES, pH 7.5, 150 mM KCl, 0.5% NP-40, 2 mM EDTA, 1 mM NaF, 0.5 mM DTT with proteinase inhibitor cocktail), and incubated on ice for 15 mins. The mitochondrial lysate was centrifuged at 12,000 rpm for 10 mins. The suspension was equally divided into two parts and incubated with 1 μg rabbit pre-immune IgG or anti-FAM210A rabbit polyclonal antibody at 4°C with rotation overnight, respectively. The protein-antibody complex was pulled down by incubation with Dynabeads Protein G (ThermoFisher Scientific) for 4 hrs and eluted using 1x SDS loading buffer. For the mass spectrometry assay, the elution was loaded onto 4-12% gradient SDS-PAGE gel and run for 10 mins. The whole lane was subjected for mass spectrometry analysis by Mass Spectrometry Resource Lab of University of Rochester Medical Center. The interaction of FAM210A and EF-Tu or ATAD3A was confirmed by Western blot following IP using indicated antibodies.
2. <u>Sample Preparation:</u> For mass spectrometry experiments, samples were run into a 4-12% SDS-PAGE gel for a short period of time to remove contaminants and create a ∼10 mm length region, allowing the total protein to be evaluated in a single gel digest. After staining with SimplyBlue SafeStain (Invitrogen), these regions were excised, cut into 1 mm cubes, de-stained, then reduced and alkylated with DTT and IAA, respectively (Sigma). Gel pieces were dehydrated with acetonitrile. Aliquots of trypsin (Promega) were reconstituted to 10 ng/μl in 50 mM ammonium bicarbonate and added so that the solution was just covering the dehydrated gel pieces. After 0.5 hr at room temperature (RT), additional ammonium bicarbonate was added until the gel pieces were completely submerged and placed at 37°C overnight. Peptides were extracted the next day by adding 0.1% TFA, 50% acetonitrile, then dried down in a CentriVap concentrator (Labconco). Peptides were desalted with homemade C18 spin columns, dried again, and reconstituted in 0.1% TFA.
3. <u>FAM210A-IP LC-MS/MS:</u> Peptides were injected onto a homemade 30 cm C18 column with 1.8 μm beads (Sepax), with an Easy nLC-1000 HPLC (ThermoFisher Scientific), connected to a Q Exactive Plus mass spectrometer (ThermoFisher Scientific). Solvent A was 0.1% formic acid in water, while solvent B was 0.1% formic acid in acetonitrile. Ions were introduced to the mass spectrometer using a Nanospray Flex source operating at 2 kV. The gradient began at 3% B and held for 2 mins, increased to 30% B over 41 mins, increased to 70% over 3 mins and held for 4 mins, then returned to 3% B in 2 mins and re-equilibrated for 8 mins, for a total run time of 60 mins. The Q Exactive Plus was operated in a data-dependent mode, with a full MS1 scan followed by 10 data-dependent MS2 scans. The full scan was done over a range of 400-1400 m/z, with a resolution of 70,000 at m/z of 200, an AGC target of 1e6, and a maximum injection time of 50 ms. Ions with a charge state between 2-5 were picked for fragmentation. The MS2 scans were performed at 17,500 resolution, with an AGC target of 5e4 and a maximum injection time of 120 ms. The isolation width was 1.5 m/z, with an offset of 0.3 m/z, and a normalized collision energy of 27. After fragmentation, ions were put on an exclusion list for 15 seconds to allow the mass spectrometer to fragment lower abundant peptides.
4. <u>Data Analysis:</u> Raw data from MS experiments were searched using the SEQUEST search engine within the Proteome Discoverer software platform, version 2.2 (ThermoFisher Scientific), using the SwissProt human database. Trypsin was selected as the enzyme allowing up to 2 missed cleavages, with an MS1 mass tolerance of 10 ppm. Samples run on the Q Exactive Plus used an MS2 mass tolerance of 25 mmu. Carbamidomethyl was set as a fixed modification, while oxidation of methionine was set as a variable modification. The Minora node was used to determine relative protein abundance between samples using the default settings. Percolator was used as the FDR calculator, filtering out peptides which had a q-value greater than 0.01.

### Lentivirus preparation and transfection

The lentivirus was produced using the 2^nd^ generation system following an established protocol^6^. Briefly, the three vectors were transfected into HEK293T cells on the first day, and collected two rounds of the supernatant at 48 hrs and 72 hrs. The supernatant was centrifuged at 2,000 g for 15 mins to remove the cell debris. The lentivirus was concentrated by centrifugation at 19,000 rpm for 2 hrs, and the pellet was dissolved in DMEM medium. For the infection, the lentivirus was added to the medium with 8 μg/ml polybrene (Santa Cruz Biotechnology) and changed back to normal cell culture medium after 24 hrs. The cells were harvested after 48 hrs and knock-down or overexpression efficiency was measured.

### RNA Northern blot analysis

RNA Northern blot analysis was performed following a previous protocol^6^. Total RNA (200 μg) was mixed with an equal volume of 2x RNA gel loading buffer (Ambion), heated at 80°C for 5 mins, ice-cooled, and run through 15% denaturing acrylamide gel electrophoresis. Total RNA was transferred to Hybond N+ (GE Healthcare) by capillary blotting using 20x SSC buffer for overnight and then fixed using a UV crosslinker. miRCURY LNA detection probes (10 pmol, Exiqon) complementary to miR-574-5p or miR-574-3p were radio-labeled with T4 polynucleotide kinase (NEB) and 1 μl [*γ*-^32^P] ATP for 1 hr at 37°C. Labeled LNA probes were heated at 95°C for 2 mins, ice-cooled, diluted in 50°C pre-warmed PerfectHyb Plus hybridization solution (Sigma) with denatured 20 μg/ml salmon sperm DNA (Ambion), and then added to the UV crosslinked pre-hybridized membrane at 50°C for 4 hrs. Membranes were washed 3 times in 2x SSC, 0.1% SDS at 50°C. The Decade Marker was used to indicate the molecular weight (Ambion). The membranes were stripped with boiled 0.1% SDS, 5 mM EDTA for 30 mins, and re-probed with radio-labeled LNA-modified U6 as a loading control.

### RIP-RT-qPCR

RIP (Ribonucleoprotein immunoprecipitation) was performed as described^4^. Protein A/G beads (40 μl) were incubated with 400 μl of homogenized heart tissue lysates (4 mg protein) for 1 h at 4°C with rotation to pre-clear. The tissue lysates was centrifuged and the supernatant collected. The rabbit anti-TDP43 (Santa Cruz Biotechnology, 2 μg) was added to the supernatant and the mixture rotated at 4°C for 1 h. The rabbit pre-immune IgG was used as a negative control antibody for IP. Protein A/G beads (40 μl) were added and incubated at 4°C for 4 h. The beads were washed 5 times with 1 ml of wash buffer (as in IP) with rotation at 4°C. Total immunoprecipitated RNA and total input RNA from 40 μl (10%) of lysate were extracted using the Trizol reagent. IPed RNA (5 μl) and 200 ng of total RNA were used in miRNA RT-qPCR assays.

### Trypan blue staining

Isolated primary adult CMs from mice were cultured in 40-mm glass dishes, treated with 10 μM of ISO for 24 hrs, and stained with Trypan blue for the assessment of cell viability of CMs. CM viability was analyzed by trypan blue dye exclusion assay. In brief, CM cultured medium was removed, and 0.04% (w/v) of trypan blue solution (VWR) was added for incubation at RT for 3-4 mins. The dead CMs appeared in blue color. CMs were visualized under the microscope. For each experiment, a total of 200 CMs were analyzed from different fields and dishes. CMs excluded from the trypan blue dye were considered as viable cells. The percentage of viability was calculated.

### Transmission electron microscopy

Murine hearts from ∼10-week old male mice (treated with vehicle or ISO injection at 30 mg/Kg/day for 4 weeks) were flushed by saline containing heparin for 10 seconds. Heart slices were cut and immediately fixed in 2.5% glutaraldehyde buffered with 0.1 M sodium cacodylate in a flat bottom tube. The fixative solution needs to be stored at 4°C and balanced to RT before use. Samples were post-fixed in 1% osmium tetroxide, dehydrated in ethanol, transitioned into propylene oxide, and then transferred into Epon/Araldite resin. Tissues were embedded into molds containing fresh resin and polymerized for 2 days at 65°C. 70 nm sections were placed onto carbon-coated Formvar slot grids and stained with aqueous uranyl acetate and lead citrate. The grids were examined using a Gatan 11 megapixel Erlansheng digital camera and Digital Micrograph software.

### Measurement of creatine kinase (CK) activity

The creatine kinase activity was measured using an ELISA based spectrophotometric Creatine Kinase Assay kit according to the manufacturer’s instructions (Abcam, Cat. No. ab155901). Isolated tissues were homogenized in the assay buffer, centrifuged at 10,000 rpm for 5 mins, and then the supernatant was used for the assay. Protein concentrations were determined by the Bradford assay. Diluted supernatants (150-200 ng) were mixed with ATP, Creatine kinase substrate, CK enzyme mix, and developer in a 96-well ELISA plate. The absorbance was read at 450 nm for 10 mins using the microplate reader. NADH was used as a standard. The CK activity was represented as U/mg/min.

### Measurement of alanine aminotransferase (ALT) activity

The alanine aminotransaminase activity was measured using the ALT assay kit (Cayman chemicals, Cat. No. 700260), based on a colorimetric method. Isolated tissues were homogenized in 100 mM Tris-HCl buffer (pH 7.8), and centrifuged at 10,000 g for 15 mins. The ALT substrate and cofactors were added to the diluted supernatant in the 96-well ELISA plate and incubated at 37 °C for 15 mins. Then the reaction was started by adding the ALT initiator and measured at 340 nm. The CK activity was represented as U/L.

### Statistical Analysis

All quantitative data were presented as mean ± SEM and analyzed using Prism 7 software (GraphPad). For comparison between 2 groups, a Student t-test was performed. For multiple comparisons among ≥3 groups, 1-way ANOVA was performed. Statistical significance was assumed at a value of P<0.05.

### Bioinformatic analysis for RNA-sequencing

Software summary:

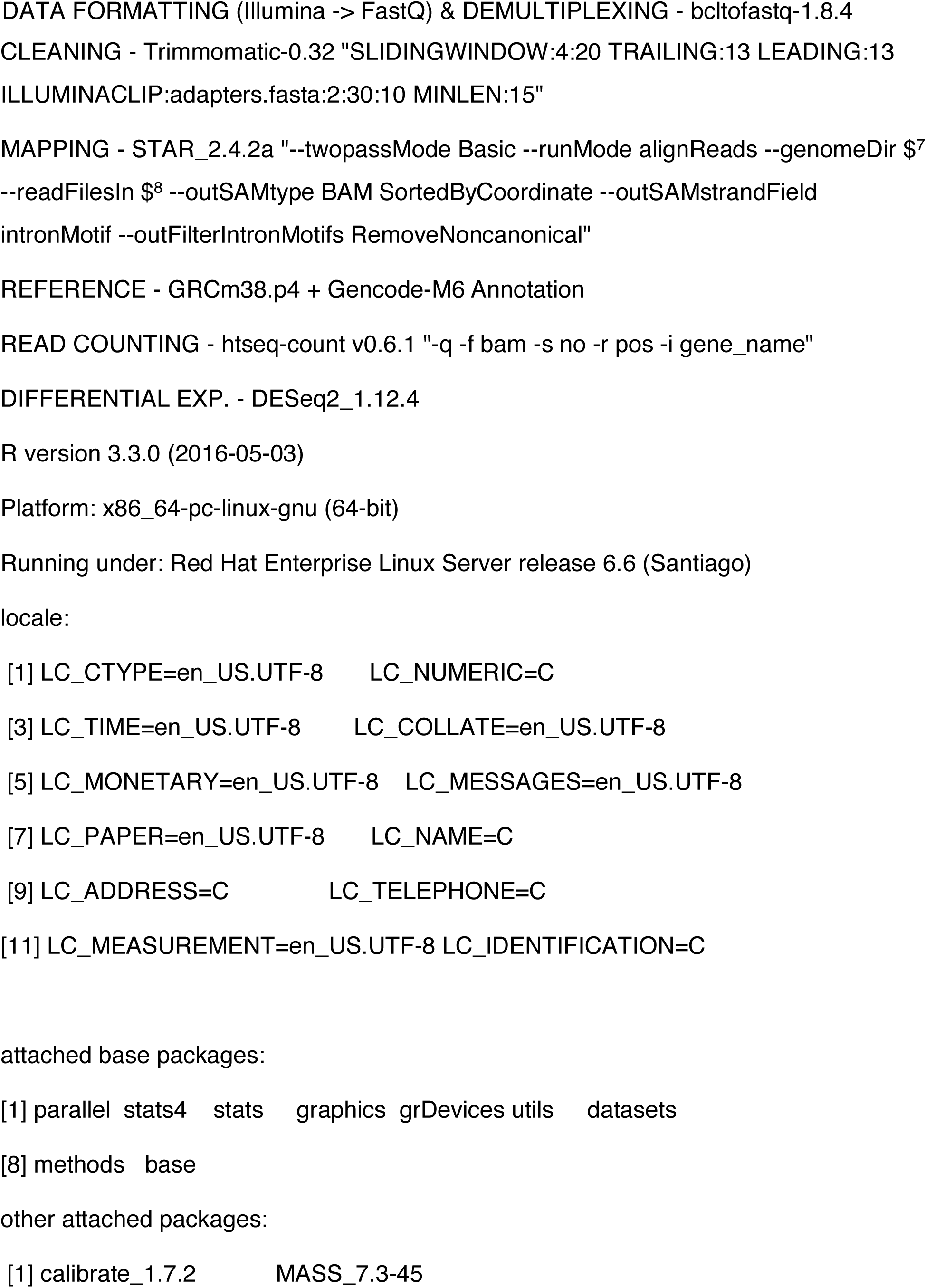
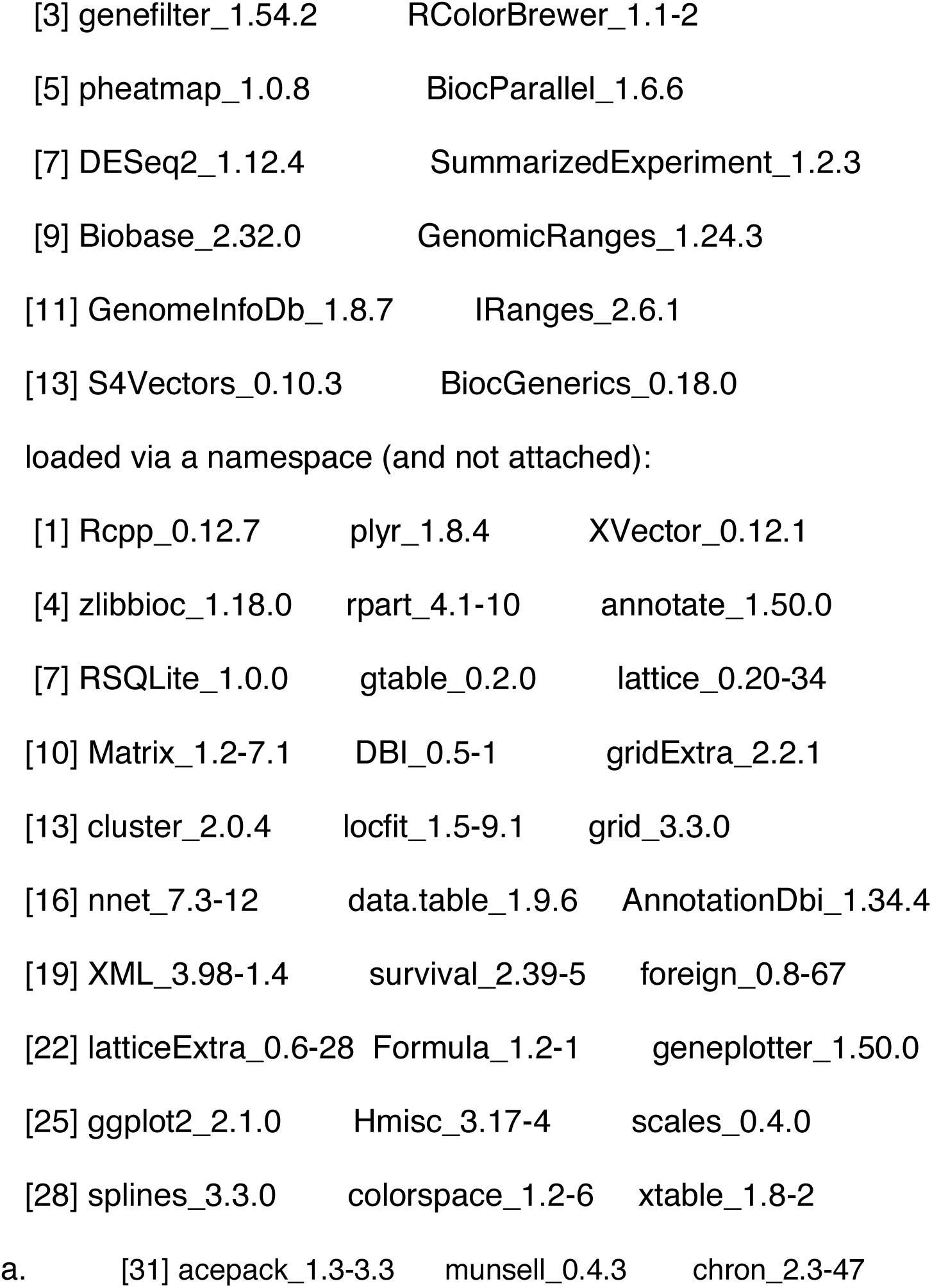

